# Non-Newtonian Blood Rheology Impacts Left Atrial Stasis in Patient-Specific Simulations

**DOI:** 10.1101/2021.06.24.449801

**Authors:** A. Gonzalo, M. García-Villalba, L. Rossini, E. Durán, D. Vigneault, P. Martínez-Legazpi, O. Flores, J. Bermejo, E. McVeigh, A. M. Kahn, J. C. del Alamo

## Abstract

The lack of mechanically effective contraction of the left atrium (LA) during atrial fibrillation (AF) disturbs blood flow, increasing the risk of thrombosis and ischemic stroke. Thrombosis is most likely in the left atrial appendage (LAA), a small narrow sac where blood is prone to stagnate. Slow flow promotes the formation of erythrocyte aggregates in the LAA, also known as rouleaux, causing viscosity gradients that are usually disregarded in patient-specific simulations. To evaluate these non-Newtonian effects, we built atrial models derived from 4D computed tomography scans of patients and carried out computational fluid dynamics simulations using the Carreau-Yasuda constitutive relation. We examined six patients, three of whom had AF and LAA thrombosis or a history of transient ischemic attacks (TIAs). We modeled the effects of hematocrit and rouleaux formation kinetics by varying the parameterization of the Carreau-Yasuda relation and modulating non-Newtonian viscosity changes based on residence time. Comparing non-Newtonian and Newtonian simulations indicates that slow flow in the LAA increases blood viscosity, altering secondary swirling flows and intensifying blood stasis. While some of these effects can be subtle when examined using instantaneous metrics like shear rate or kinetic energy, they are manifested in the blood residence time, which accumulates over multiple heartbeats. Our data also reveal that LAA blood stasis worsens when hematocrit increases, offering a potential new mechanism for the clinically reported correlation between hematocrit and stroke incidence. In summary, we submit that hematocrit-dependent non-Newtonian blood rheology should be considered in calculating patient-specific blood stasis indices by computational fluid dynamics.

## 1 Introduction

Ischemic stroke is a leading cause of mortality and disability. Ischemic strokes account for two thirds of all strokes, which are estimated to be 26 million per year worldwide [1]. Approximately 30% of these ischemic strokes are directly associated with atrial fibrillation (AF), a common arrhythmia affecting 35 million people worldwide [2]. An additional 30% of ischemic strokes are categorized as embolic strokes from undetermined source (ESUS), and increasing evidence suggests many of these also have a cardioembolic origin [3]. Specifically, a significant fraction of these ESUS may be linked to left atrial thrombosis, both in the presence of subclinical AF and in sinus rhythm [4]. Therefore, cardioembolism is a major source of stroke. However, the indication of anticoagulant drugs must be carefully balanced against the risk of bleeding. In patients with AF, clinical tools to estimate stroke risk are based on demographic and comorbidity factors (*e.g*., the CHA_2_DS_2_-VASc score), which do not capture patient-specific thrombogenesis mechanisms and have a modest predictive accuracy [5, 6]. Likewise, indiscriminate anticoagulation of ESUS patients has not proven effective in preventing recurrent strokes despite increasing bleeding event rates [7]. In summary, because left atrial thrombosis is a major source of ischemic strokes, precision patient-specific indices of intracardiac thrombosis risk may improve current indications of anticoagulant therapy for stroke prevention.

Thrombosis risk increases with the concurrence of three phenomena: endothelial damage or dysfunction, the presence of procoagulatory factors in the blood, and increased blood stasis [8]. In the left atrium (LA), most thrombi form inside the left atrial appendage (LAA), a narrow hooked sac that accounts for approximately 5-10% of LA volume and which varies greatly in morphology among individuals [9]. The irregular beating of the heart with a lack of organized atrial contraction, together with anatomical factors promotes the appearance of LAA stasis [10, 11]. Intracavitary blood stasis and thrombogenesis risk can be quantified through blood residence time, *i.e*., the time spent by blood particles inside a cardiac chamber [12], whose calculation requires detailed knowledge of the flow velocity field. Progress in computational fluid dynamics (CFD) algorithms and computer processing capacity now make it feasible to simulate blood flow in four-chamber domains [13], two-chamber domains (typically the left heart) [14, 15, 16], and single-chamber domains (typically the left ventricle) [17, 18, 19, 20, 21]. Some of these simulations include models of heart valve motion [19, 14, 16, 22] and some can resolve the intricate domain surface caused by endocardial trabeculation [22]. Motivated by the need for patient-specific prediction of LAA blood stasis, the number of CFD simulations of LA flow has increased rapidly in the last few years [23].

Blood inside the cardiac chambers is usually modeled as a Newtonian fluid based on the assumption that its shear rate is high enough to prevent the formation of red blood cell (RBC) aggregates and that, even if some low-shear pockets may occasionally form, their lifetime is shorter than the RBC aggregation timescale [24, 25]. While an exhaustive validation of these modeling assumptions can be challenging, Newtonian CFD simulations in chambers other than the LAA provide intracardiac flow velocity fields that agree reasonably well with phase-contrast MRI data from the same patients [18, 26]. Nevertheless, calculating residence time inside the LAA has two peculiarities that make it particularly sensitive to non-Newtonian effects. First, CFD data suggest that both low shear rate and high residence time coexist inside the LAA [27, 28], making this site prone to RBC aggregation. Consonantly, spontaneous echocardiographic contrast or “smoke”, which is caused by red blood cell aggregates under low shear [29], is frequently observed in the LAA by transesophageal echocardiography and is associated with blood stasis [30]. Second, the residence time reflects the aggregate effects of flow transport over multiple heartbeats in contrast to other flow metrics like kinetic energy or shear stress which are not accumulative. Therefore, slight differences in instantaneous velocity could gradually accumulate to generate appreciable differences in residence time. Considering that blood is a shear-thinning fluid, we hypothesize that Newtonian CFD analysis could underestimate blood stasis, and this effect could be more severe for regions of significant stasis.

The present study evaluates whether blood’s shear-thinning rheology significantly affects CFD estimations of LAA blood stasis. To this end, we carried out numerical simulations of left atrial flow considering a Carreau-Yasuda constitutive relation and compared the results with those from Newtonian simulations [28]. We examined six patients covering a wide range of atrial morphologies and functions, three of whom had LAA thrombosis or a history of transient ischemic attacks (TIAs). Our results indicate that blood stasis increases in low-shear regions when non-Newtonian rheology is considered, especially in the LAA. This trend is robust for different constitutive laws that consider the timescale of RBC aggregation and the hematocrit value, i.e., the volume fraction occupied by RBCs. Finally, our simulations indicate that LAA blood stasis and, by implication, LAA thrombosis risk, increase with the hematocrit level both in healthy and AF subjects. These data insinuate a novel mechanism for the clinically observed connection between high hematocrit and stroke, which had been previously attributed solely to alterations in cerebral blood flow [31].

## 2 Methods

### 2.1 Patient population, image acquisition, image reconstruction, and mesh generation

We retrospectively considered the six left atrial models as in our previous work [28], consisting of 4D (3D, time-resolved) patient-specific segmentations of human left atria obtained from computed tomography (CT) imaging. Three patients had normal LA volume and function, were imaged in sinus rhythm, and did not have a left atrial appendage (LAA) thrombus. The other three patients were imaged in atrial fibrillation (AF), had an enlarged LA chamber, and impaired LA global function. Two of these subjects had an LAA thrombus which were removed digitally before running the simulations, and one had a history of transient brain ischemic attacks (TIAs). Digital removal of the thrombi was performed by comparing the primary dynamic acquisition with a late contrast acquisition obtained 22 seconds after the primary one. Cardiac-gated cine CT scans were performed following standard clinical protocols at each participating center (NIH, Bethesda, MD and UCSD, La Jolla, CA). The images were reconstructed using the CT scanner manufacturers’ standard algorithms, yielding DICOM files with resolutions between 0.32 mm and 0.48 mm in the *x*-*y* axial plane and 0.5 mm to 1 mm in the *z*-direction. Time-resolved images were obtained at regularly spaced time points across the cardiac cycle, ranging between 5% and 10% of the R-R interval.

The computational LA meshes were generated in four steps using ITK-SNAP [32] and MATLAB. The first step consisted of segmenting the 3D LA anatomy 4D CT time frame and identifying the pulmonary veins (PVs) inlets, mitral valve (MV) annulus, and LAA. For each 3D segmentation, a triangular surface mesh was created and then resampled to match the computational fluid dynamics (CFD) solver’s resolution [33]. The resulting triangular meshes were registered across the cardiac cycle and their positions, yielding a coherent triangle vertex and centroid cloud. The resulting point clouds were registered using the coherent point drift registration algorithm implemented in the open-access software toolbox developed by Myronenko and Song [34]. Specifically, we used the non-rigid low-rank kernel approximation. Finally, the positions of these points were expressed as a Fourier temporal series to provide interpolated boundary conditions to the CFD solver at time points not contained in the 4D CT sequence. Figure 1 and Table 1 summarize the main anatomical and functional features of the patients’ atria. More details on patient selection, imaging acquisition and reconstruction, and mesh generation can be found elsewhere [28].

**Figure 1:**
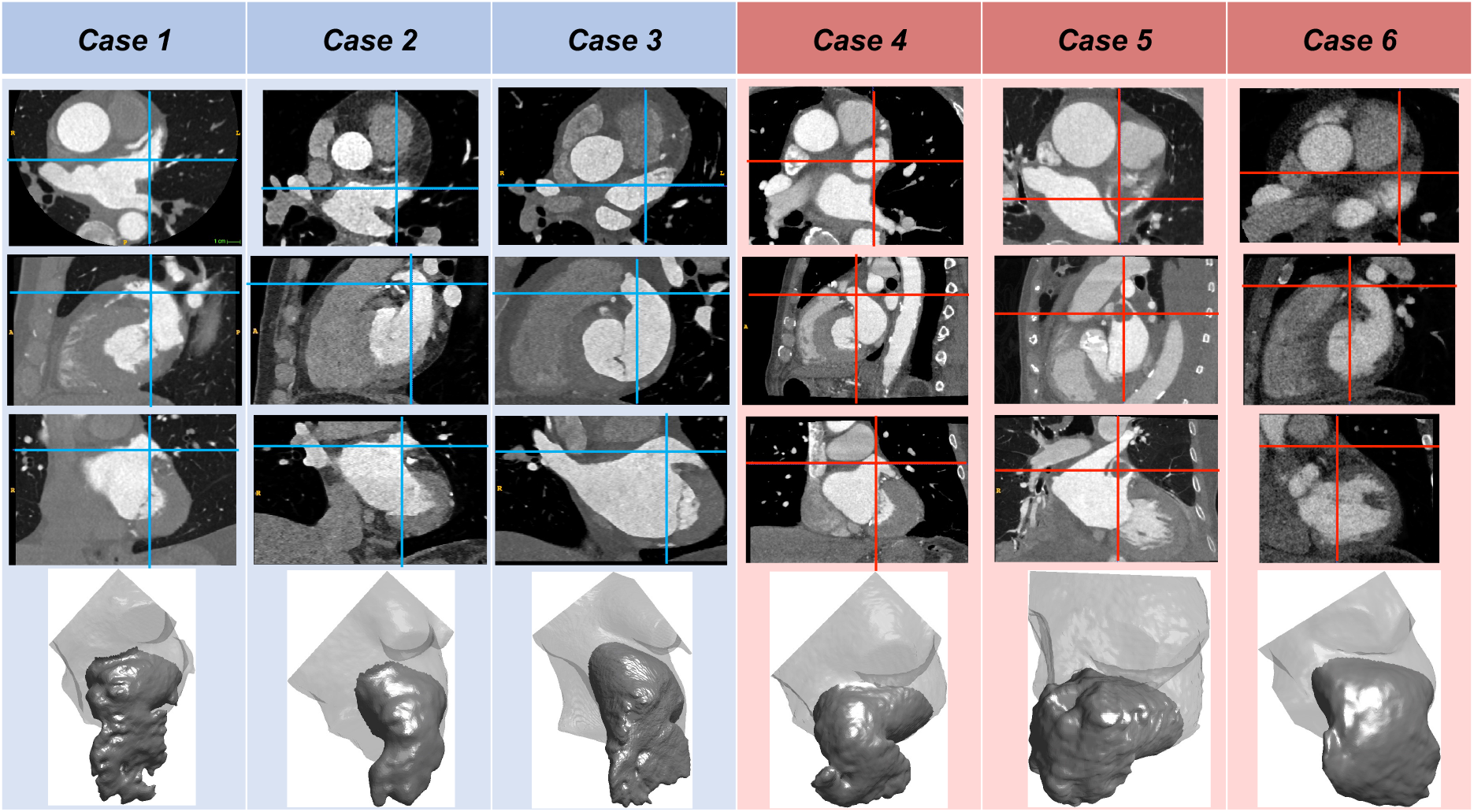
Anatomical Features of Patient-Specific Simulation Cases. Computerized tomography (CT) images (axial, sagittal, and coronal plane sections, rows 1-3 respectively) and segmented appendages (row 4) of the six left atria considered in this study. The images show left atrial appendage (LAA) thrombi in cases 5 and 6. The vertical and horizontal lines in each CT view indicate the intersections with the other two view planes. The images correspond to an instant at 50% of the R-R interval.

**Table 1:** Summary of anatomical and functional parameters of the left atrium (LA) and its appendage (LAA) in the LAA thrombus/transient ischemic attack-negative (left three columns) and LAA thrombus/transient ischemic attack-positive (right three columns) groups. Mean volume indicates time-averaged volume. Ostium aspect ratio (AR) is defined as the ratio between the length of the ostium major (*L_Major_*) and minor (*L_Minor_*) axes. Ostium AR and ostium Area shown are values time-averaged over the cardiac cycle. Reynolds number at the mitral valve annulus (Re_MV_) is computed using a characteristic diameter 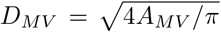, where *A_MV_* is the area of the MV, the peak velocity during the E-wake 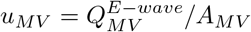, and the Newtonian blood viscosity *ν*_∞_. EF, ejection fraction; PVs, blood volume that enters the LA during LV systole; PVd, blood volume that enters the LA during LV diastole; PVa, blood volume that exits the LA due to reverse flow volume through the pulmonary veins during atrial contraction; E/A ratio, ratio of peak mitral velocities during early diastole (E-wave) and atrial contraction (A-wave).

| Subject Number | 1 | 2 | 3 | 4 | 5 | 6 |
| --- | --- | --- | --- | --- | --- | --- |
| **LAA Thrombus (or history of TIA)** | No | No | No | TIA's | Yes | Yes |
| **Atrial Fibrillation** | No | No | No | Persistent | Persistent | Persistent |
| **Sinus Rhythm** | Yes | Yes | Yes | No | No | No |
| **Mean LA Volume (ml)** | 86.6 | 70.1 | 115 | 145 | 157 | 180 |
| **Min. LA Volume ($V_{LA}^{min}$, ml)** | 59.6 | 49.0 | 87.2 | 119 | 150 | 157 |
| **Max. LA Volume ($V_{LA}^{max}$, ml)** | 108 | 91.2 | 145 | 155 | 165 | 205 |
| **Mean LAA Volume (ml)** | 6.94 | 4.85 | 14.3 | 10.7 | 15.5 | 22.0 |
| **Min. LAA Volume ($V_{LAA}^{min}$, ml)** | 4.32 | 3.14 | 10.2 | 9.10 | 13.8 | 19.8 |
| **Max. LAA Volume ($V_{LAA}^{max}$, ml)** | 8.97 | 6.28 | 17.9 | 11.6 | 17.4 | 24.7 |
| **Ostium Area ($mm^2$)** | 4.17 | 4.01 | 7.65 | 7.22 | 7.29 | 8.42 |
| **Ostium AR ($L_{Major}/L_{Minor}$)** | 1.53 | 1.39 | 1.49 | 1.45 | 1.45 | 1.83 |
| **Pre-A-wave Volume ($V_{LA}^{pre-A}$, ml)** | 94.3 | 72.3 | 119 | 149 | 153 | 178 |
| **LA Sphericity** | 0.80 | 0.82 | 0.78 | 0.86 | 0.83 | 0.81 |
| **Global (LA EF) $\left[ \frac{V_{LA}^{max} - V_{LA}^{min}}{V_{LA}^{max}} \right]$** | 0.45 | 0.46 | 0.40 | 0.23 | 0.091 | 0.23 |
| **LAA EF $\left[ \frac{V_{LAA}^{max} - V_{LAA}^{min}}{V_{LAA}^{max}} \right]$** | 0.52 | 0.50 | 0.43 | 0.22 | 0.21 | 0.20 |
| **Expansion Index $\left[ \frac{V_{LA}^{max} - V_{LA}^{min}}{V_{LA}^{min}} \right]$** | 0.81 | 0.86 | 0.66 | 0.30 | 0.10 | 0.31 |
| **Passive EF $\left[ \frac{V_{LA}^{max} - V_{LA}^{pre-A}}{V_{LA}^{max}} \right]$** | 0.13 | 0.21 | 0.18 | 0.039 | 0.073 | 0.13 |
| **Booster EF $\left[ \frac{V_{LA}^{pre-A} - V_{LA}^{min}}{V_{LA}^{pre-A}} \right]$** | 0.37 | 0.32 | 0.27 | 0.20 | 0.020 | 0.12 |
| **Conduit Volume / LV Stroke Volume $\left[ 1 - \frac{V_{LA}^{max} - V_{LA}^{min}}{SV} \right]$** | 0.33 | 0.50 | 0.41 | 0.46 | 0.80 | 0.63 |
| **PVs (ml)** | 43 | 38 | 51 | 36 | 17 | 49 |
| **PVd (ml)** | 65 | 58 | 60 | 52 | 63 | 94 |
| **PVs-PVd ratio** | 0.66 | 0.66 | 0.84 | 0.69 | 0.27 | 0.52 |
| **PVa (ml)** | 17 | 7.3 | 12 | 13 | 0.14 | 1.4 |
| **PVa duration (cycles)** | 0.11 | 0.11 | 0.13 | 0.12 | 0.039 | 0.068 |
| **A-wave duration (cycles)** | 0.22 | 0.25 | 0.30 | 0.18 | 0.21 | 0.23 |
| **PVa – A-wave duration ratio** | 0.49 | 0.43 | 0.44 | 0.64 | 0.18 | 0.30 |
| **E/A ratio** | 1.64 | 2.36 | 3.03 | 2.05 | 3.86 | 3.65 |
| **Re<sub>MV</sub> $\left[ Re_{MV} = \frac{u_{MV} D_{MV}}{\nu_{\infty}} \right]$** | 4297 | 6266 | 6315 | 4361 | 4111 | 8058 |

### 2.2 Computational Fluid Dynamic

We used a modified version of the CFD code TUCAN [35], which we had previously adapted to simulate incompressible, Newtonian flow in the left atrium [28]. To account for non-Newtonian effects, we consider the Navier-Stokes equations

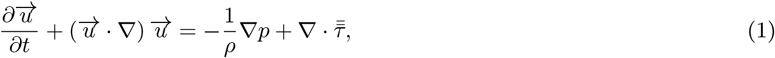

where 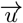 and *p* are respectively the velocity and pressure fields, *ρ* is the fluid density, and *τ* is the stress tensor. The latter is defined as 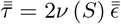, where *ν* is the fluid kinematic viscosity, 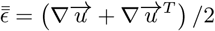 is the rate-of-strain tensor, 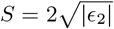 is the shear rate, and *ϵ_2_* the second invariant of 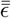. The constitutive relation between *ν* and *S* is based on the Carreau-Yasuda model [36], both in its original form and with modifications to account for blood’s thixotropic behavior under unsteady flow. Section 2.3 below provides more details about the constitutive models implemented in our CFD code. The interested reader can find validation examples of the Newtonian and non-Newtonian versions of TUCAN in [37] and in the Supporting Information, respectively.

The Navier-Stokes equations were integrated in time by a low-storage, three-stage, semi-implicit Runge–Kutta scheme using a fractional-step method. The constitutive law was treated semi-implicitly by splitting the kinematic viscosity coefficient into two terms, *i.e., ν* (*S*) = *ν_imp_*+Δ*ν_exp_*(*S*). In this decomposition, the implicit part, *ν_imp_*, is a constant coefficient and the explicit part, Δ*ν_exp_*(*S*), can vary in space and time and is non-linear in the velocity field through its dependence on the shear rate. Consequently, the viscous stresses were split as 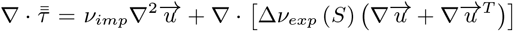. The first, linear term was integrated in time implicitly leading to a linear system of equations each time-step, like in the Newtonian case. The second, non-linear term was integrated explicitly to avoid non-linear systems of equations every temporal iteration. Because the implicit treatment of the viscous terms is unconditionally stable while the explicit treatment is not, it is advisable to choose a constant value of *ν_imp_* that minimizes the amplitude of the explicit viscous terms. We chose *ν_imp_* = *ν*_∞_ (i.e., the Newtonian viscosity) but note that the implementation is independent of this particular choice, offering some control over the stability of the temporal integration.

The time step Δ*t* of each simulation was constant and set to keep the Courant-Friedrichs-Lewy (*CFL*) number below 0.2 across the whole simulation. To account for the additional explicit terms originating from Δ*ν_exp_*(*S*), the Courant number was re-defined as 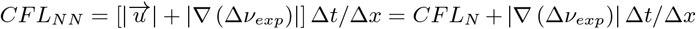, where the subindices *N* and *NN* stand for Newtonian and non-Newtonian. The spatial discretization was performed on a staggered Cartesian grid with a centered, second-order, finite difference scheme. The grid spacing, Δ*x* = 0.51 mm (256^3^ grid points), was isotropic and uniform. We initialized each non-Newtonian simulation using the results of [28], taking the converged velocity field and residence time computed using Newtonian rheology and nominal resolution for each subject. The flow variables were interpolated to a coarse resolution grid (Δ*x_c_* = 16Δ*x*/9, 144^3^ grid points), and then ran for 15 heartbeats at 60 beats per minute (bpm) to let the flow transition from Newtonian to non-Newtonian. The motivation for this strategy was to save computational resources as the coarse resolution grid requires (144/256)^4^ = 0.1 times the number of floating-point operations per cardiac cycle (a factor of 144 per spatial direction and one more factor due to Δt through the *CFL* condition). Finally, we interpolated the flow variables into the 256^3^ nominal grid and ran the simulations for 6 more heartbeats at 60 bpm. To evaluate the convergence, we computed the cycle-to-cycle difference of the velocity magnitude, which was smaller than 2% during the last 3 cycles for all the cases presented here.

The time spent by blood particles inside the LA chamber was denoted residence time, *T_R_*, and calculated together with the velocity field by solving the forced transport equation [38]

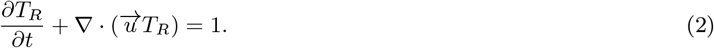

This equation was integrated in time explicitly with a low-storage, three-stage Runge-Kutta scheme. Its spatial discretization was a third-order WENO scheme to balance accuracy with preventing spurious oscillations caused by the Gibbs phenomenon in regions with sharp gradients [39]. Together with the shear rate, the residence time is used to indicate blood stasis and non-Newtonian rheology.

The flow was driven by the heart’s wall motion obtained from patient-specific 4D CT images, which was imposed using the immersed boundary method (IBM) as previously described [28]. In short, the segmented LA surface was placed inside a 13-cm cubic Cartesian mesh, and free-slip boundary conditions were imposed at the cube boundaries. No-slip boundary conditions were imposed at the LA inner surface by adding an appropriate volumetric force term (i.e., the IBM force) to the Navier-Stokes equations. To enforce flow rates through the PVs inlets, a non-cylindrical buffer region of immersed boundary points was defined by extending each PV inlet plane section 12 grid points upstream. A volumetric force (akin to the IBM force) was applied in this region to smoothly accelerate the flow to 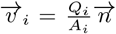, where *Q_i_*, *A_i_* and 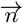 are the flow rate, inlet area, and normal vector of the *i-th* PV inlet (for *i* = 1…4). The *Q_i_*’s were determined from mass conservation as 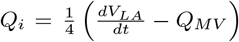, where *V_LA_* is the time-dependent left atrial volume, 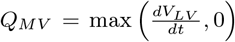 is the flow through the mitral valve, and *V_LV_* is the time-dependent left ventricular volume. Both *V_LV_* and *V_LA_* are obtained from each patient’s 4D CT scan. The ¼ factor evenly splits the total flow rate through the four PVs. The boundary conditions at the mitral annulus outlet are imposed on the plane section at the downstream end of the atrial segmentation, which moves inside the cubic simulation domain as the LA walls deform. When the mitral valve is closed, the mesh points in that section are treated as a Lagrangian no-slip immersed boundary similar to the rest of the atrial wall. When the mitral valve is open (i.e., *Q_MV_* > 0), no boundary conditions are imposed on the mitral lid of the segmentation.

Finally, since eq. 2 only contains first-order derivatives, the only boundary condition needed for the residence time is *T_R_* = 0 at the PV inlets. This condition was enforced with a similar procedure as the one used for the velocity, defining an appropriate volumetric force in the buffer region upstream of the PVs inlets. More details about the boundary conditions of the simulations can be found in [28].

### 2.3 Non-Newtonian constitutive models

We ran simulations using two different rheological models. Each model was parameterized considering two hematocrit values *Hct*, leading to four constitutive laws in the simulations. First, we used the original Carreau-Yasuda model

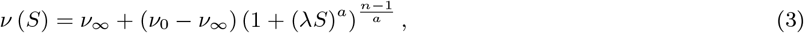

where, *λ*, *n* < 1, and *a* are blood hematocrit-dependent parameters and *ν*_∞_ is the reference Newtonian viscosity corresponding to the limit *S* ≫ *λ*^-1^. In this model, yield-stress is represented by prescribing a large value to the zero-shear-rate viscosity coefficient, *ν*_0_, which we prescribed as *ν*_0_ = 16*ν*_∞_ similar to De Vita *et al*.[40].

To investigate the effect of *Hct*, we considered two different sets of parameter values derived from the literature, namely λ = 8.2 s, *a* = 0.64, *n* = 0.2128 from Leuprecht and Perktold [41], and λ = 3.313 s, *a* = 2.0, n = 0.3568 from Al-Azawy *et al*.[42]. Although these studies considered *ν*_∞_ = 0.035 cm^2^/s, we chose *ν*_∞_ = 0.04 cm^2^/s to compare with our previous Newtonian simulations [28], which where run with a constant kinematic viscosity equal to 0.04 cm^2^/s. Figure 2A shows that the shear-rate-dependent kinematic viscosities obtained from the two parameter sets differ significantly. We used the blood rheology experiments of Chien *et al*. [43] to estimate their associated *Hct* values. Specifically, we least-squares fitted *Hct* in eq. 3 to match the viscosity empirical law of Chien *et al*. (see eq. I in ref. [43]), achieving best fits for *Hct* = 37.4 and 55.0 (Figure 2A). Hence, we refer to these models as CY-37 and CY-55 throughout our manuscript. Blood hematocrit levels can vary significantly with age and sex, with all-age normal ranges (defined as the 2.5-97.5 percentile range) being [34–36,48–50] for females and [39–40, 54] for males [44, 45]. Thus, the CY-37 and CY-55 models are helpful to estimate the smallest and largest non-Newtonian effects to be expected in LA hemodynamics.

**Figure 2:**
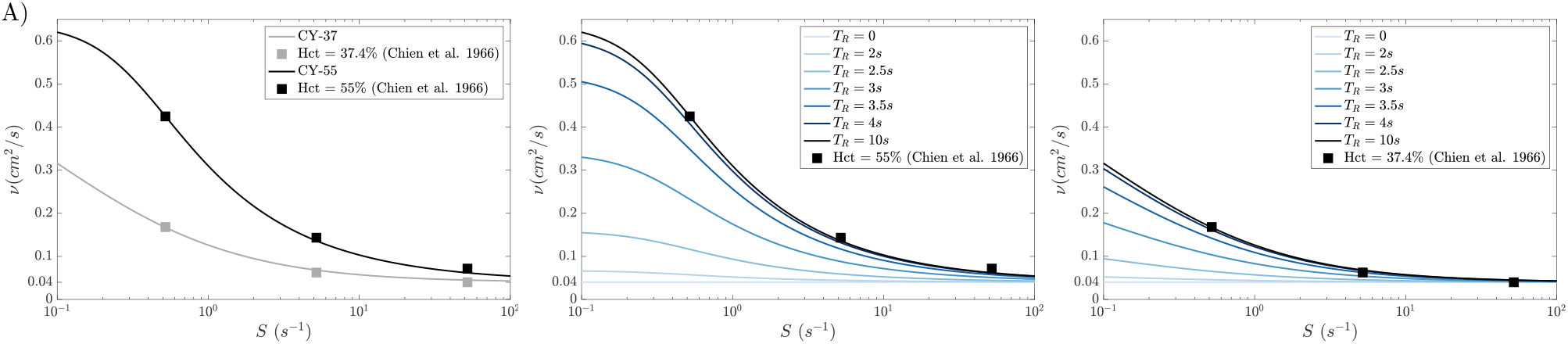
Non-Newtonian constitutive laws used in the present simulations. **A)** Kinematic viscosity *ν* (*S*) as a function of shear rate *S* for the two versions of the Carreau-Yasuda law (eq. 3) used in our simulations. Grey line: CY-37 model with λ = 8.2 s, *a* = 0.64, *n* = 0.2128, *ν*_0_ = 16*ν*_∞_, and *ν*_∞_ = 0.04 cm^2^/s; black line: CY-55 model with λ= 3.313 s, *a =* 2.0, n = 0.3568, *v_0_=* 16v∞, and v∞= 0.04 cm^2^/s. The square symbols correspond to datapoints from the empirical law of Chien *et al*.[43] for hematocrit values *Hct*= 37.4% (grey) and Hct = 55% (black). Kinematic viscosity *ν* (*S*, *T_R_*) as a function of shear rate *S* and residence time *T_R_* for the residence-time-activated Carreau-Yasuda model **B)** CY-T_R_-55 and **C)** CY-T_R_-37 (eqs. 4–5). Black squares come from Chien *et al*.’s [43] data for the respective *Hct* values.

The Carreau-Yasuda constitutive relation assumes microstructural equilibrium, but blood is a thixotropic fluid that exhibits time-dependent rheology because the formation and rupture of RBC rouleaux does not occur infinitely fast [36]. While the observation of LA and LAA echocardiographic “smoke” in patients demonstrates that rouleaux do form in these chambers [29, 30], the unsteady nature of atrial hemodynamics calls for evaluating the interplay between fluidic and thixotropic timescales. Huang and Fabisiak [46] modeled the thixotropic behavior of blood using a power-law term restrained by an exponential term that considered the cumulative exposure of RBCs to shear, i.e., proportional to 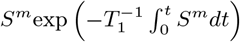. In this model, *T*_1_ is the time scale associated to rouleaux formation and breakdown. Parameter fitting to human blood samples in healthy subjects yielded *T*_1_ values ranging between 3.6 and 6.2 seconds. Consonantly, Schmid-Schönbein *et al*. [47] had previously reported the half-time for RBC aggregation in blood to range between 1.2–6.0 seconds (mean 3.5 seconds) for normal blood samples and between 0.4–3.0 seconds for hypercoagulable samples (mean 1.13, 1.69, andn 0.94 seconds in myeloma patients, diabetics, and pregnant women at term, respectively). More recently, Arzani [25] proposed a modified Carreau-Yasuda constitutive relation where non-Newtonian effects are activated based on the local value of the residence time. In the same spirit, we ran simulations where blood viscosity was given by

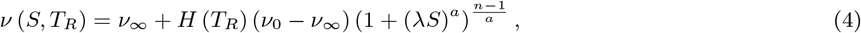

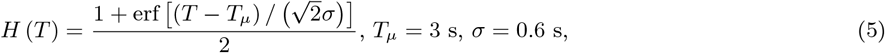

where *T_μ_* and *σ* are the mean and standard deviation of the error function used for the activation time of non-Newtonian effects. The function *H* (*T*) provides a smooth transition from Newtonian to non-Newtonian behavior for residence times comparable to experimentally reported RBC aggregation timescales. We used *T_μ_* = 3 s consistent with the literature consensus for normal patients [47, 46], noting that this choice likely provides a conservative estimate of non-Newtonian effects since hypercoagulable samples have lower values of *T_μ_*, described above, which are less restrictive to rouleaux formation. We ran simulations at *Hct* = 37 and 55 with the constitutive model in eqs. (4–5), which we denoted as CY-*T_R_*-37 and CY-*T_R_*-55. Figure 2B illustrates the dependence of the CY-*T_R_*-55 model with residence time, showing that non-Newtonian effects are negligible for *T_R_* ≲ 2 s, gradually increase for 2 s ≲ *T_R_* ≲ 10 s, and become indistinguishable from those of the classic Carreau-Yasuda model for *T_R_* ≳ 10 s. The dependence of the CY-*T_R_*-37 model with residence time is similar (Figure 2C).

### 2.4 Quantification of non-Newtonian effects

To compare the velocity fields from Newtonian and non-Newtonian simulations, we defined the orientation (*ϵ_O_*) and normalized magnitude (*ϵ_M_*) non-Newtonian velocity differences as:

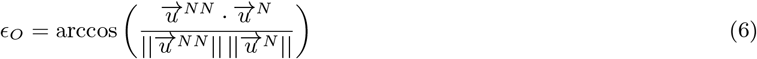

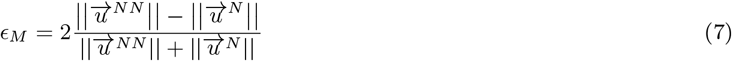

where 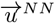 and 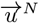 are the velocity field of the Non-Newtonian and Newtonian simulations, respectively. The orientation error *ϵ_O_* measures the angle between 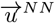 and 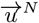. Positive values of correspond to higher magnitude of the velocity on the non-Newtonian case than in the Newtonian case.

To evaluate the regions where non-Newtonian effects are important, we computed the ratio between the non-Newtonian and Newtonian viscosities, *ν*/*ν*_∞_. To quantify how non-Newtonian rheology influenced blood stasis in the LAA we used two different metrics. First, the change in the residence time defined as:

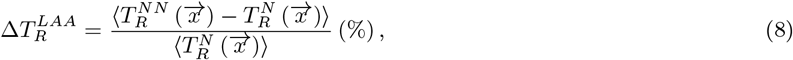

where the super index LAA indicates averaging in space inside the LAA, and the brackets 〈 〉 indicate averaging in time over the last three cardiac cycles. Second, we focused on the distal LAA, the region with the highest residence time where thrombi are most likely to form. Specifically, we considered the 33% voxels inside the LAA most distant from the ostium and computed the percentage of those occupied by stagnant blood, defined as *T_R_* larger than several cutoff values (6, 8, and 10 s). These values are summarized in Table 2.

**Table 2:**
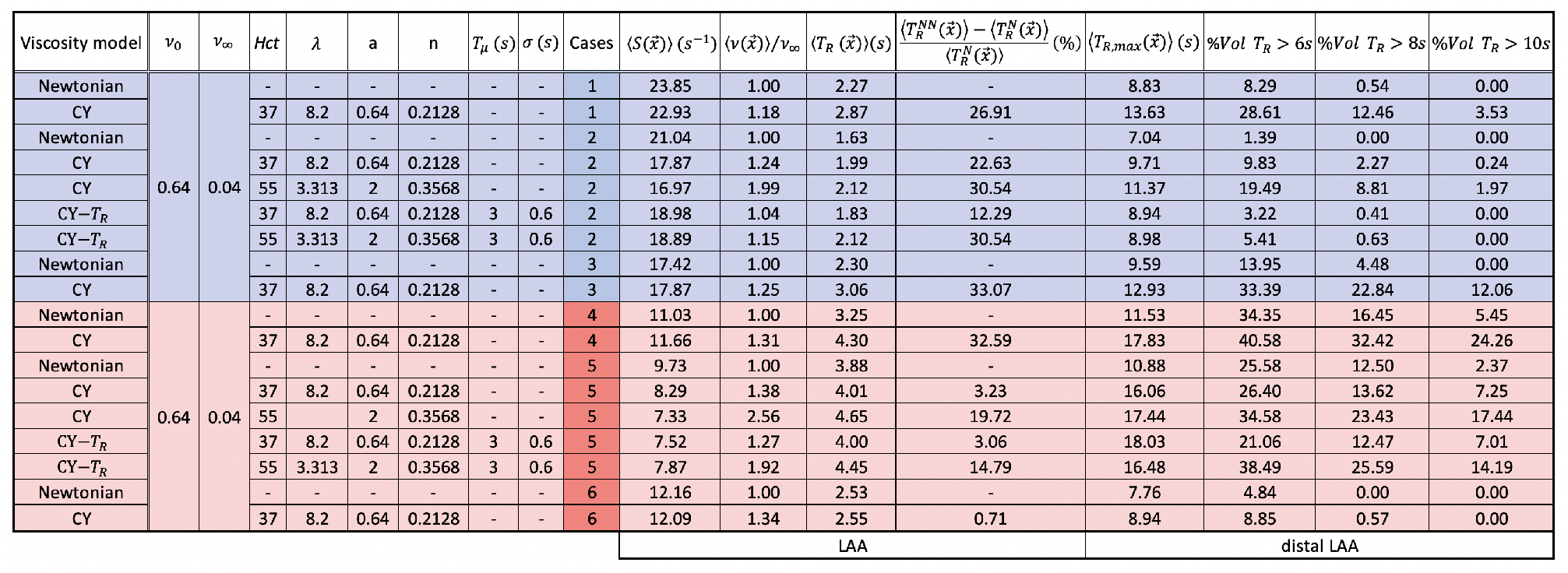
Summary of blood rheology models and cases studied, including time-averaged hemodynamical statistics evaluated in the left atrial appendage (LAA) and the distal LAA. *Hct*, hematocrit level; Carreau-Yasuda model parameters (*ν*_0_, *λ*_∞_, *a* and *n*), and modified Carreau-Yasuda model additional parameters (*T_μ_* and *σ*), see section 2.3 for description; %*V ol T_R_* > *τ*, percentage of volume with residence time larger than the time threshold *τ*. Metrics between 〈 〉 are time averages over the last three cycles highlighted with a grey shaded region in Figures 6 and 7.

## 3 Results

### 3.1 Flow visualization in Newtonian and non-Newtonian left atrial simulations

We first evaluated whether the LA body and LAA flow patterns were affected by non-Newtonian blood rheology. Figure 3 displays 3D maps of blood velocity vectors in a representative normal subject (case 2), computed by the Newtonian simulations as well as the non-Newtonian CY-37, and CY-55 simulations. The vectors near the left PVs, the right PVs, the LAA, and the mitral annulus are colored differently to facilitate visualization. Magnified 3D vector maps inside the LAA are also displayed. To quantify how non-Newtonian effects modified the magnitude and orientation of blood velocity, the figure also includes volumetric maps of *ϵ_O_* and *ϵ_M_* in the LAA (defined in eqs. 6 and 7 above). Snapshots corresponding to LA diastole, LV early filling, and LA systole are represented. For reference, the figure includes flow rate profiles of the pulmonary veins (PVs) and mitral (MV) annulus, together with LA and LAA chamber volume vs. time. Figure 4 shows similar simulation results for a representative AF, LAA-thrombus-positive patient (case 5). Figures SI3-SI 6 of the Supporting information show analogous plots for the other 4 simulated patients.

**Figure 3:**
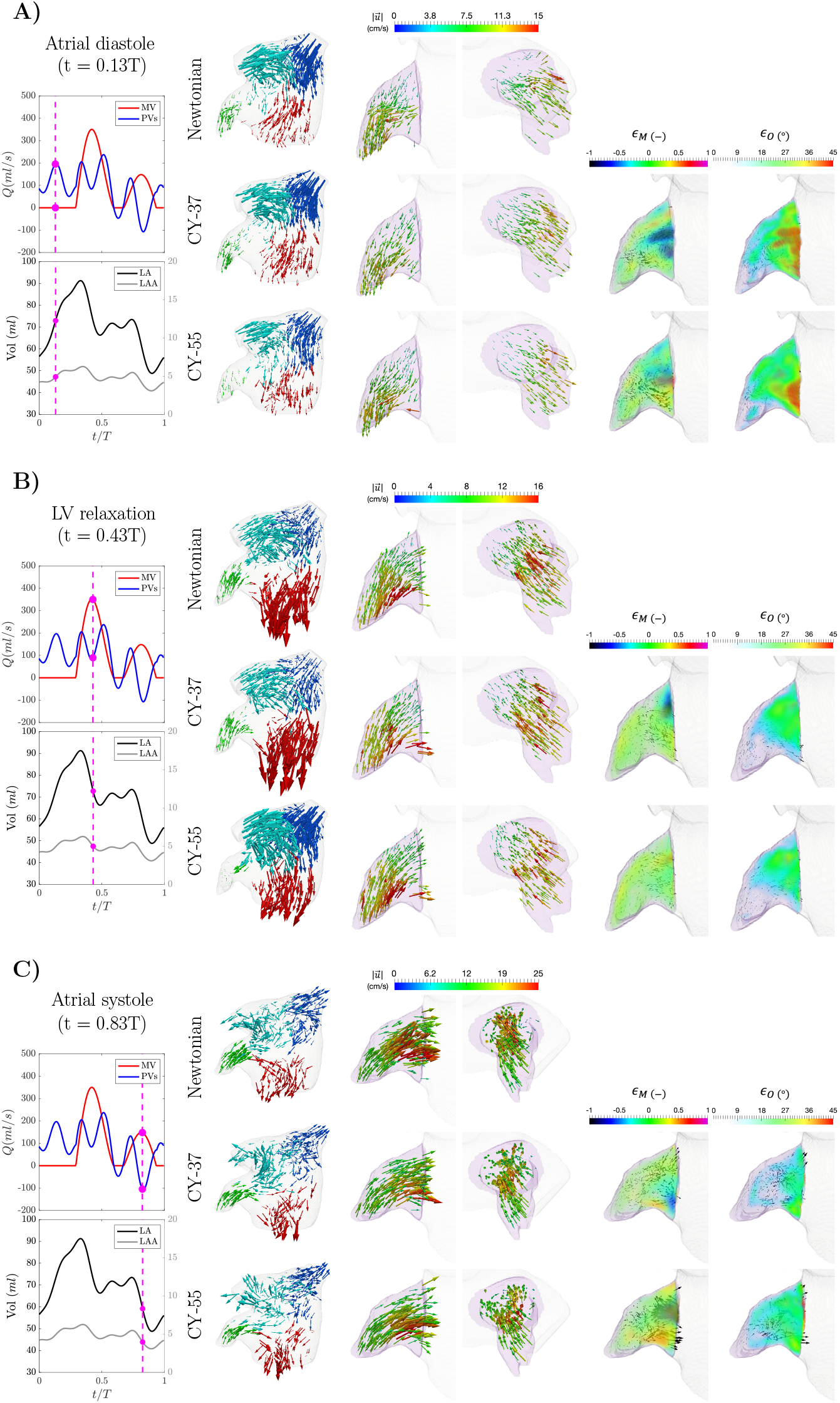
Flow visualization of left atrial and left atrial appendage (LAA) hemodynamics from Newtonian and non-Newtonian simulations. Case2: Subject with normal atrial function and no LAA thrombus. Each panel displays a snapshot of the 3-D blood flow velocity in the whole left atrium (1^st^ column), two amplified views of the LAA in different orientations (2^nd^ and 3^rd^ columns), and two LAA views showing the differences in velocity magnitude, (4^th^ column) and orientation, *ϵ_O_*, (5^th^column) between Newtonian and non-Newtonian flow. In the 1^st^column, the vectors are colored according to their proximity to the right pulmonary veins (blue), left pulmonary veins (cyan), left atrial appendage (green) and mitral valve (red). In the 2^nd^and 3^rd^columns, the vectors are colored according to the velocity magnitude. The vectors in all views are scaled with the velocity magnitude. The colors of the 4^th^ and 5^th^ columns are transparent for *ϵ_M_* = 0 and for *ϵ_O_*, 5°, respectively. Each panel also includes time histories of the flow rate through the mitral valve (red) and the cumulative flow rate through the pulmonary veins (blue), and of the volumes of the left atrium (black) and the left atrial appendage (grey). The magenta bullets indicate the instant of time represented in the vector plots of each panel. The flow vectors inside the left atrium are represented at three instants of time. **A)** Atrial diastole and peak flow rate through the pulmonary veins (t = 0.13 s). **B)** Left ventricular diastole and peak flow rate through the mitral valve (E-wave, t = 0.43 s). **C)** Atrial systole and peak backflow rate through the pulmonary veins (t = 0.83 s).

**Figure 4:**
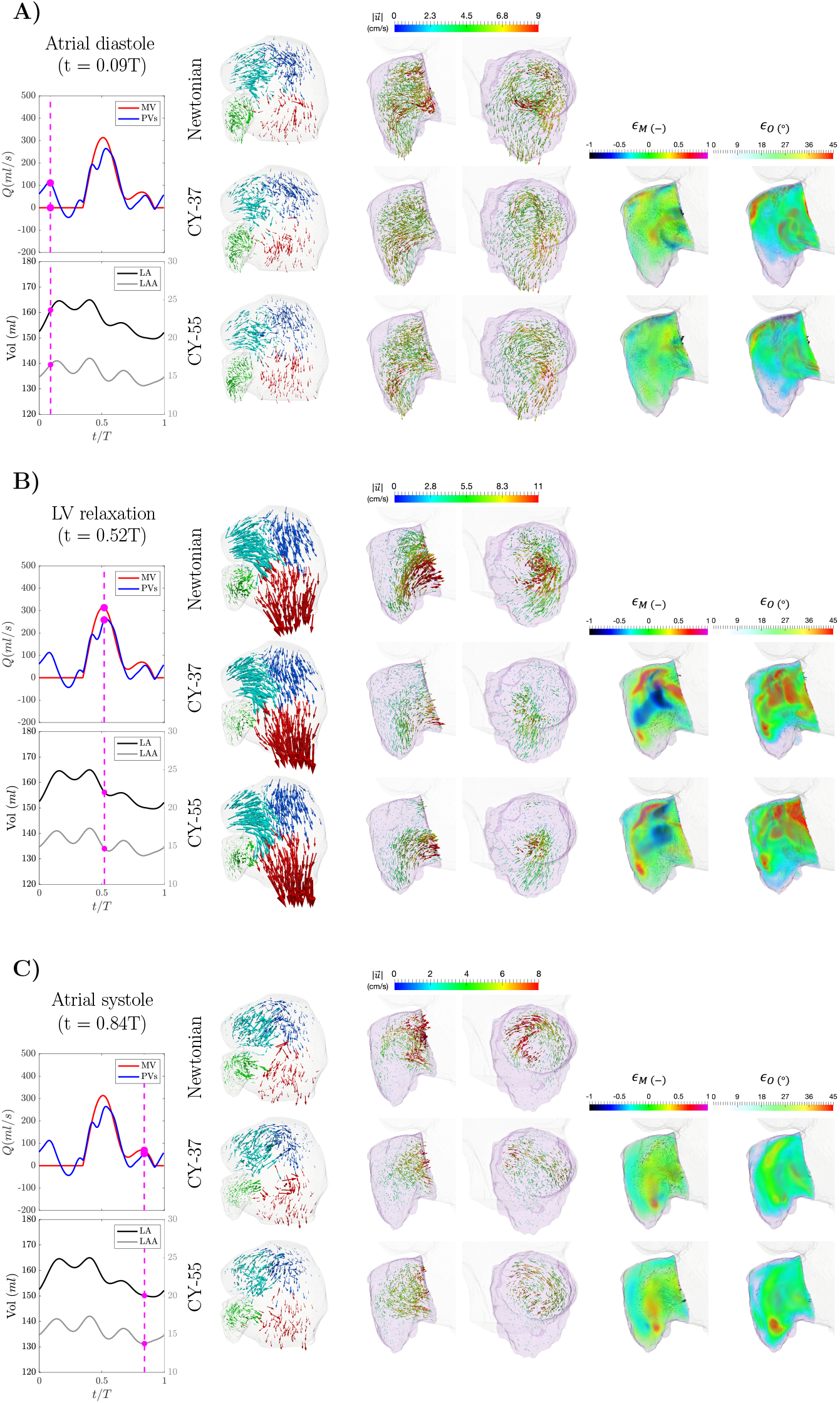
Flow visualization of left atrial and left atrial appendage (LAA) hemodynamics from Newtonian and non-Newtonian simulations. Case 5: Atrial fibrillation patient with LAA thrombus (digitally removed before running the simulations). Vector maps of the 3-D blood flow velocity in the whole left atrium (1^st^ column), two amplified views of the LAA in different orientations (2^nd^ and *3^rd^* columns), and two LAA views showing the differences in velocity magnitude (4^th^column) and orientation (5^th^ column) between Newtonian and non-Newtonian flow, using the same format as Figure 3. **A)** Atrial diastole and peak flow rate through the pulmonary veins (t = 0.09 s). **B)** Left ventricular diastole and peak flow rate through the mitral valve (E-wave, t = 0.52 s). **C)** Atrial systole and peak backflow rate through the pulmonary veins (t = 0.84 s).

For the most part, the large-scale flow patterns inside the LA body were unaltered in the non-Newtonian CY-37 and CY-55 simulations, as compared to the Newtonian simulations. In the healthy cases, atrial chamber expansion during atrial diastole drove a strong flow from the PVs to the mitral annulus (Figure 3A), which was sustained by ventricular suction and elastic atrial recoil during LV filling (Figure 3B). Atrial contraction (Figure 3C) kept squeezing blood out of the LAA and the LA body into the ventricle, also creating backflow from the LA into the PVs. Non-Newtonian effects did not seem to significantly alter the large-scale flow patterns in the LA body of AF patients either. The LA mainly acted as a conduit with strong fluid motion from the PVs into the ventricle during LV early filling (Figure 4B), while the flow was practically arrested during the remainder of the cardiac cycle (Figure 4A, C).

On the other hand, we found appreciable qualitative and quantitative differences in flow patterns inside the LAA. First, the filling and drainage LAA jets were altered when non-Newtonian effects were considered. In the healthy subject (case 2), the non-Newtonian changes in velocity orientation and magnitude, c_o_and *_M_*, were significant in proximal LAA inflow and outflow jets, near the ostium. These changes were most appreciable during atrial diastole and systole (3A and C). In the LAA-thrombus-positive AF patient (case 5), the flow patterns in the LAA exhibited considerable secondary swirling flows, probably associated with the more voluminous and tortuous LAA shape of this patient (e.g., 15.5 ml vs. 4.85 ml, see Table 1). These swirling motions showed significant non-Newtonian changes, both in velocity magnitude and orientation in the whole LAA volume (Figure 4). Overall, the time-volume-averaged values of *ϵ_O_* and *ϵ_M_* (Table SI1) were appreciable and comparable across all non-Newtonian simulations.

### 3.2 Blood inside the left atrial appendage continuously experiences low shear rates consistent with non-Newtonian rheology

Since low shear rates and RBC aggregation are hallmarks of non-Newtonian blood rheology, we analyzed the shear rate 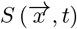 in our simulations. Figure 5 shows instantaneous snapshots of 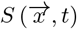 from normal case 3, an AF patient with decreased LA reservoir function (case 4, history of TIAs), and an AF patient with reduced reservoir and booster functions (case 5, thrombus positive). To visualize LA vortices in the atrial body, we also represented iso-surfaces of the second invariant of the velocity gradient tensor (*Q* = 1000 s^-2^). The data come from CY-37 simulations, which were available for the three patient-specific geometries.

**Figure 5:**
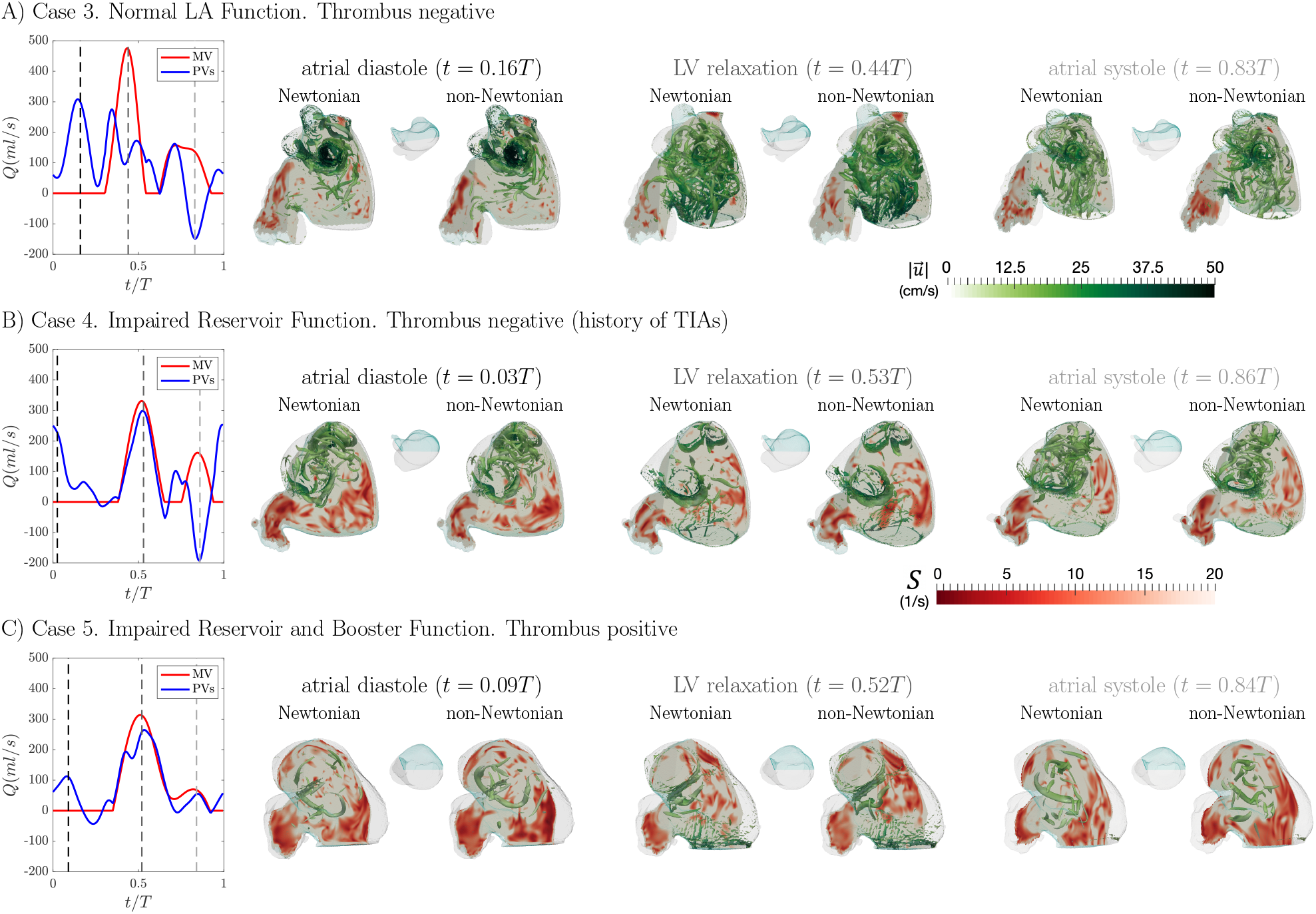
Left atrial shear rate and vortex structures inside left atrium (LA) of subjects with or without left atrial appendage (LAA) thrombus and with different atrial function. Three subjects are shown, one without thrombus and normal atrial function (**panel A**), one with a history of transient ischemic attacks and impaired reservoir function (**panel B**), and one with an LAA thrombus and both impaired reservoir and booster functions (**panel C**). Each panel includes plots of the time histories of flow rate through the mitral valve (red) and the cumulative flow rate through the pulmonary veins (blue). For each subject, instantaneous snapshots of the shear rate S are shown in two oblique plane sections of the LAA and the left atrial body, indicated the sagittal view insets. Additionally, the vortex patterns inside the LA are represented using an iso-surface of the second invariant of the velocity gradient tensor (*Q* = 1000 s^-2^), and colored according to the local velocity magnitude. Snapshots from both Newtonian and CY-37 simulations (Carreau-Yasuda model of eq. 3, hematocrit value Hct = 37) are included. These snapshots correspond to three instants of time: **1)** atrial diastole (i.e., peak flow rate through the pulmonary veins), **2)** left ventricular rapid filling (i.e., the E-wave, peak flow rate through the mitral valve), and **3)** atrial systole (i.e., the A-wave of left ventricular filling and peak backflow rate through the pulmonary veins). These three instants of time are indicated with dashed vertical lines in the flow-rate plots at the left-hand-side of each panel.

Given the relatively large Reynolds number and the unsteady nature of LA flow, the shear rate fluctuated in space and time. However, despite these fluctuations, 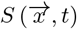 remained low (< 10 s^−1^) in large portions of the LAA across the whole cardiac cycle. Both in the Newtonian and non-Newtonian simulations, these LAA shear rates are low enough to trigger non-Newtonian effects, since the CY-37 and CY-55 constitutive models yield *ν*/*ν*_∞_ = 1.3 and 2.0 respectively at *S* = 10 s^-1^(Figure 2). In addition, for the non-Newtonian cases, the increased viscosity in the LAA yields slightly lower shear rates (see Table 7), except for case 6. The subjects with decreased atrial function had particularly low shear rates inside their LAA. These subjects also displayed low *S* regions inside the atrial body, but these regions were not sustained over long times. The distributions of 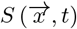 for the Newtonian and CY-37 blood simulations were similar in that they produced low shear rates inside the LAA across the whole cardiac cycle. The LA vortex structures were qualitatively similar, although there was a trend for non-Newtonian simulations to yield increased velocities along intense vortex cores, which is to be expected in shear-thinning fluids.

### 3.3 Blood viscosity in Newtonian and non-Newtonian left atrial simulations

To evaluate the variations in atrial blood viscosity caused by the shear rate dependence and thixotropy of blood rheology, we determined the normalized time-averaged kinematic viscosity over the last three heartbeats of our simulations, 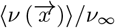. We mapped this quantity in two oblique intersecting plane sections of the LA body and the LAA for the same two cases considered in Figures 3 and 4 (Figure 6A). In addition, we plotted time histories of the median of *ν*/*ν*_∞_ inside the LA body and LAA (Figure 6B), and the probability density functions of *ν*/*ν*_∞_ inside the LAA for the last simulated cycle (Figure 6C). Consistent with the sustained presence of low-shear regions inside the left atrium, Figure 6 indicates that the CY-37 and CY-55 simulations yielded higher viscosity than the Newtonian simulation. This increase was particularly appreciable inside the LAA.

**Figure 6:**
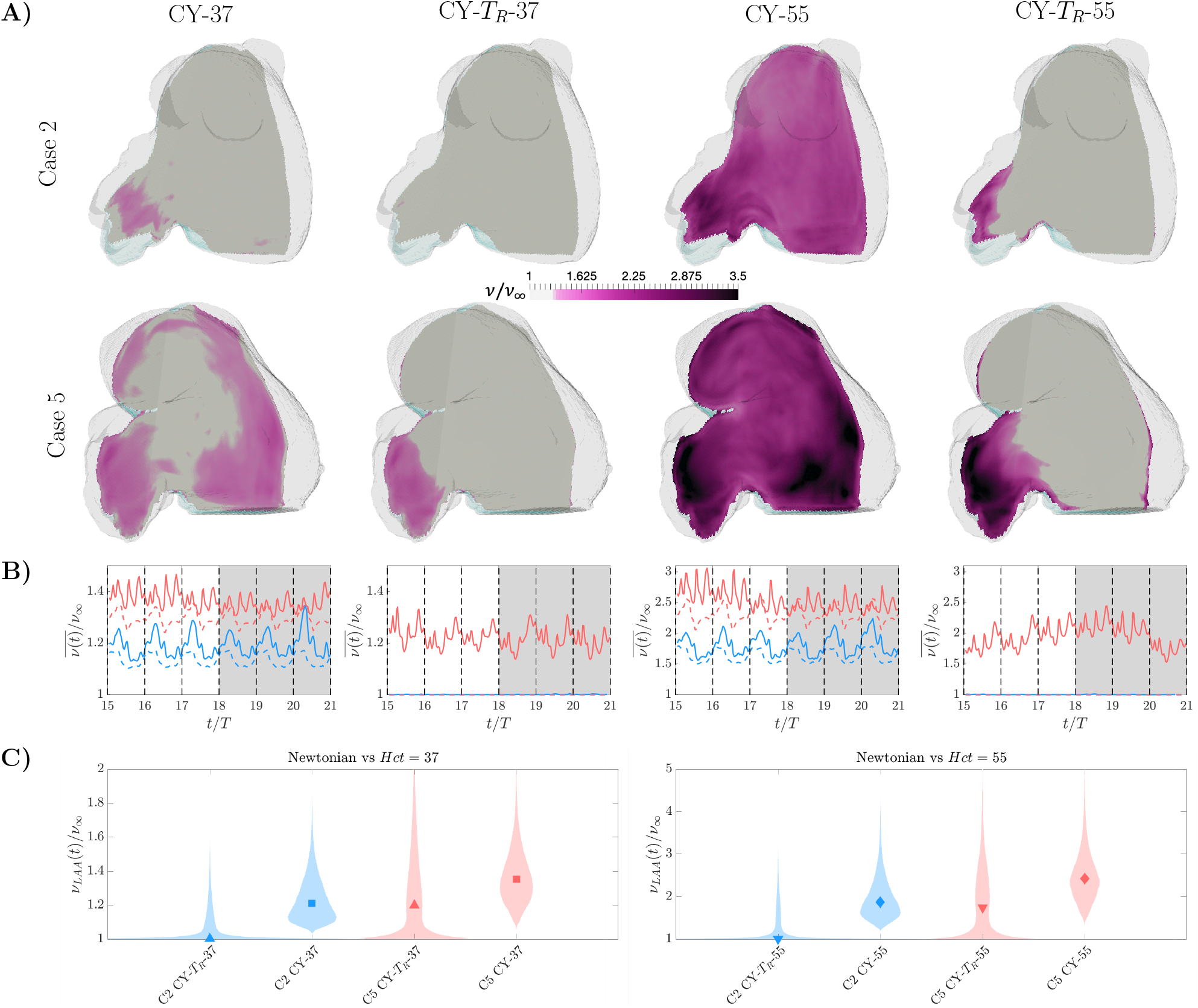
Left atrial viscosity for non-Newtonian different constitutive laws. **A)** Spatial distribution of kinematic viscosity. Two subjects are shown, one without thrombus and normal atrial function (**top**), and one with a left atrial appendage (LAA) thrombus (digitally removed before running the simulations) and both impaired reservoir and booster functions (**bottom**). For each subject, the time-averaged, normalized viscosity 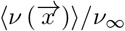 is shown in two oblique plane sections of the LAA and the left atrial (LA) body, similar to Figure 5. Data from simulations with low-hematocrit CY-37 (1^st^ column), low-hematocrit, residence-time-activated CY-*T_R_*-37 (2^nd^ column), high-hematocrit CY-55 (3^rd^ column), and high-hematocrit, residence-time-activated CY-*T_R_*-55 (4^th^ column) constitutive laws are included. **B)** Time histories of kinematic viscosity from the healthy, thrombus-negative subject (blue) and the thrombus-positive patient (red) with AF shown in **panel A**. Data from the last 6 cycles of the simulations are plotted, with the grey shaded region indicating the three cycles used to calculate time averages (i.e., 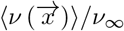 in **panel A** and **panel C**). Lines represent the median. Solid lines indicate the LAA while dashed lines indicate the LA body. **C)** Violin plots of the probability density function (shaded patches) inside the LAA, together with the median (symbol). Colors as in **panel B**.

Table 2 displays statistics of non-Newtonian effects for all our simulations. The low-hematocrit CY-37 simulations produced moderate increases in viscosity for healthy subjects (range 1.18-1.25), which were mostly confined inside the LAA. There was a trend for the CY-37 simulations to produce higher increases in viscosity in AF patients (range 1.31-1.38) than in healthy ones, with more extensive zones of elevated 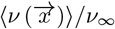 inside the LAA and and the LA body.

The high-hematocrit CY-55 simulations produced significantly larger viscosity values all over the LA body and LAA, with 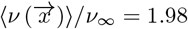 and 2.56 for the healthy and AF patients.

Previous Newtonian CFD analyses (reviewed in [23]) and our own [28] suggest that LA blood residence time can be comparable to the timescale associated with blood’s thixotropic behavior [47, 46]. Thus, the CY-55 and CY-37 models, which assume steady-state blood rheology, could overestimate blood viscosity in low residence time zones. To investigate how the competition between RBC aggregation kinetics and flow unsteadiness influences the results of the CFD analysis, we ran the two patient-specific cases of Figures 3 and 4 using the low CY-*T_R_*-37 and high-hematocrit CY-*T_R_*-55 constitutive laws (eqs. 4–5). In these thixotropic constitutive relations, non-Newtonian effects only kick in if the local residence time is longer than the activation time *T_μ_* = 3 s. Consistent with this *T_R_*-dependent activation, the CY-*T_R_*-55 model yielded viscosities that were very close to the constant Newtonian reference value inside the whole LA body and in the proximal part of the LAA (Figure 6). In contrast, the CY-55 model produced significant non-Newtonian effects across the whole LA body, with 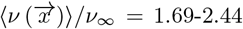. The same trend was observed when comparing the CY-*T_R_*-37 and CY-37 simulations, although including *T_R_*-dependent activation had less dramatic results at *Hct* = 37 than at Hct = 55 because non-Newtonian effects are moderate at low hematocrit. The next section characterizes how the viscosity variations caused by the constitutive laws we investigated (*i.e*. CY-37, CY-55, CY-T_R_-37, and CY-T_R_-55) affect the predictions of blood stasis from the CFD analysis.

### 3.4 Blood stasis in Newtonian and non-Newtonian left atrial simulations

We examined how non-Newtonian blood rheology affects LAA blood stasis by mapping the blood residence time obtained with the CY-37, CY-55 constitutive laws, and comparing it to previous results reported using the constant viscosity assumption [28]. The *T_R_* maps indicate that most of the LA body’s blood pool is cleared and replenished every 1-2 cardiac cycles (*T_R_* ≤ 2 s, grey shades in Figure 7A). On the other hand, the LAA residence time can reach large values, particularly in the LAA regions most distal from the ostium (Figure 7A). The increase in residence time between the LA body and the appendage was larger in patients with impaired atrial function (*e.g*., case 5 in Figure 7), whose LAA contracted and expanded weaker than in healthy subjects (see LA functional parameters in Table 1). The LAA was also the site where the residence time maps produced by non-Newtonian simulations differed most from the Newtonian ones. These results are consistent with our findings that non-Newtonian effects were most significant inside the LAA (Figure 6) and that these effects weakened the transport of blood inside this chamber (Figures 3 and 4). The CY-37 and CY-55 simulations exacerbated high-TR stagnant regions by augmenting their size and maximum residence time. Conversely, regions of moderate or low *T_R_* were not affected as much. These differences were significant even in the low hematocrit simulation, CY-37, and became more pronounced in the high hematocrit case, CY-55. For instance, the apical LAA residence times predicted by the CY-37 simulation for the healthy and AF cases of Figure 7 were 32% and 64% higher than their Newtonian values, respectively, whereas these differences were 54% and 67% when considering the CY-55 constitutive law (see Table 2).

**Figure 7:**
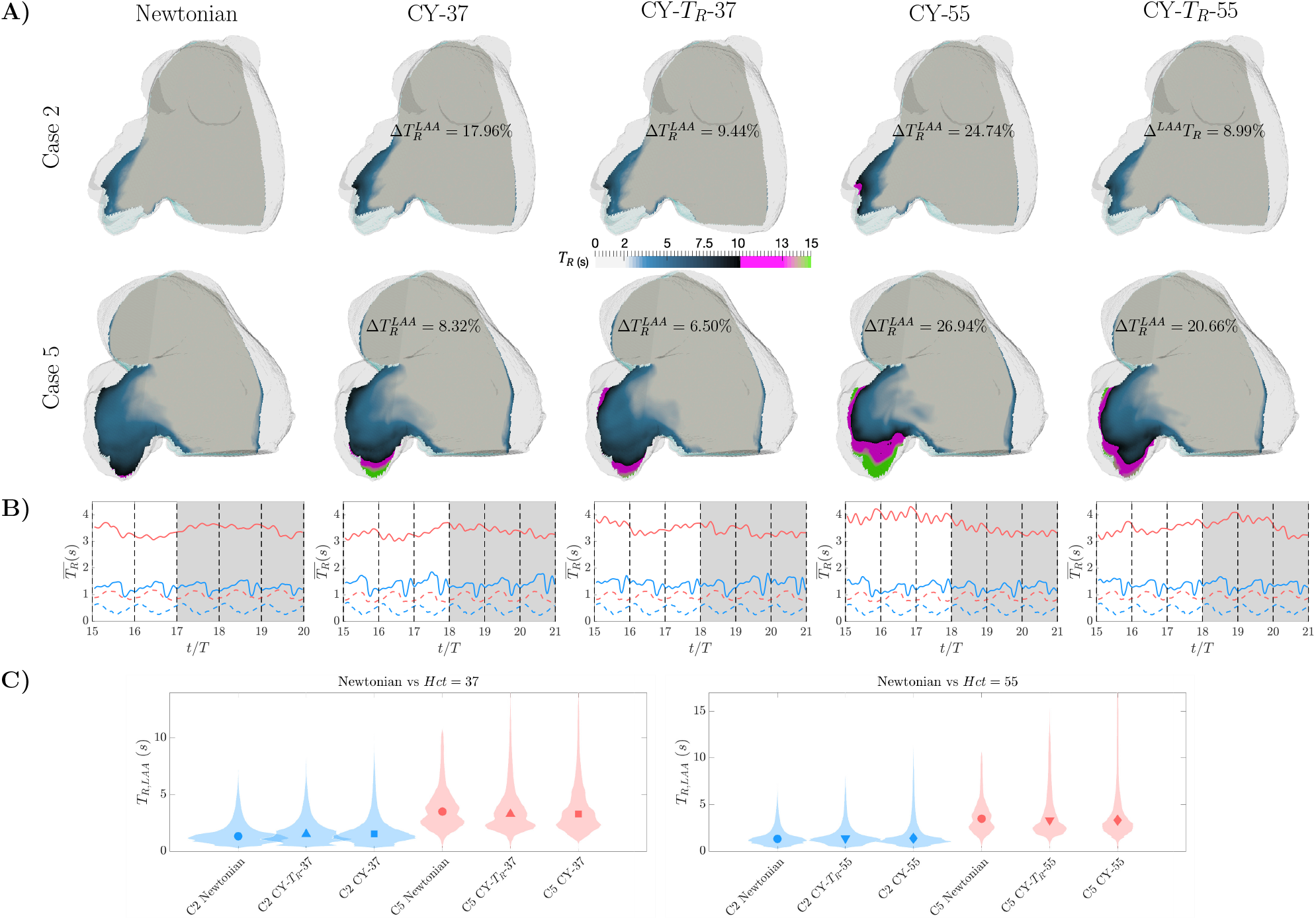
Left atrial blood residence time for Newtonian and non-Newtonian different constitutive laws. Panels of this Figure are presented in the same format as Figure 6, but including Newtonian results in the left column. **A)** Spatial distribution of blood residence time. **B)** Time histories of blood residence time inside the left atrial body and the left atrial appendage (LAA). **C)** Violin plots of the probability density function inside the LAA.

As expected, the thixotropic non-Newtonian simulations (CY-*T_R_*-37, CY-*T_R_*-55) predicted lower LAA residence time values than their equilibrium counterparts (CY-37, CY-55), albeit still larger than the Newtonian simulations. However, the situation was different in the LA body, where all non-Newtonian and Newtonian simulations yielded similar results for *T_R_*. Although, this result might seem counter-intuitive, below we argue that it is the direct consequence of residence time being dictated by mass conservation in the LA body (see Discussion section).

Figure 7A also shows that the fraction of the distal LAA predicted to be stagnant increased notably when including non-Newtonian effects. Table 2 shows that, when comparing the low-*Hct* model CY-37 with the reference Newtonian simulations, the stagnant blood volume in the distal LAA increased from 9% to 26% on average for healthy subjects and from 22% to 30% for AF patients for moderate cutoff values (6 s), whereas it increased from 0.1% to 6.6% in healthy subjects and from 2.9% to 11% in AF patients for high cutoff values (10 s). Likewise, the maximum residence time increased from 9 to 13 s on average for healthy subjects and from 10 to 15 s for AF patients. In addition, both the stagnant distal LAA volume and the maximum residence time increased with *Hct* for a given given constitutive law (*i.e*., CY-37 vs. CY-55 and CY-T_R_-37 vs. CY-T_R_-55). Overall, these results suggest that neglecting non-Newtonian rheology can lead to underestimating the size of stagnant regions inside the LAA as well as their level of stasis.

Two other metrics commonly used in CFD analyses to summarize cardiac hemodynamics and, in particular, stasis are the flow mean shear rate *S* and kinetic energy, 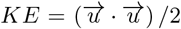, respectively. Figure 8 displays the median together with the *25^th^* and *75^th^* percentiles of these two variables inside the LAA. The data are phase-averaged for the last three cycles of the simulations and dislayed vs. time for heartbeat. The data come from the same normal subject (case 2) and AF patient with an LAA thrombus (case 5) considered in the previous figures. Simulations results from the constitutive relations CY-37, CY-55, CY-*T_R_*-37, and CY-*T_R_*-55 are plotted together with those from the reference Newtonian simulations (grey). In both subjects, non-Newtonian effects moderately shifted the distributions of *S* and *KE* towards lower values of these variables. In the healthy subject, the shift was most pronounced during the atrial diastole and LV relaxation, while it was barely detectable during atrial systole (being less noticeable in *KE*, see green arrows). In the AF patient, the shift towards lower *S* and *KE* was sustained throughout the whole cardiac cycle. The contrast between the large increase of *T_R_* and the moderate differences in *S* or *KE* supports the idea that velocity changes accumulating over multiple heartbeats significantly impact residence time and blood stasis.

**Figure 8:**
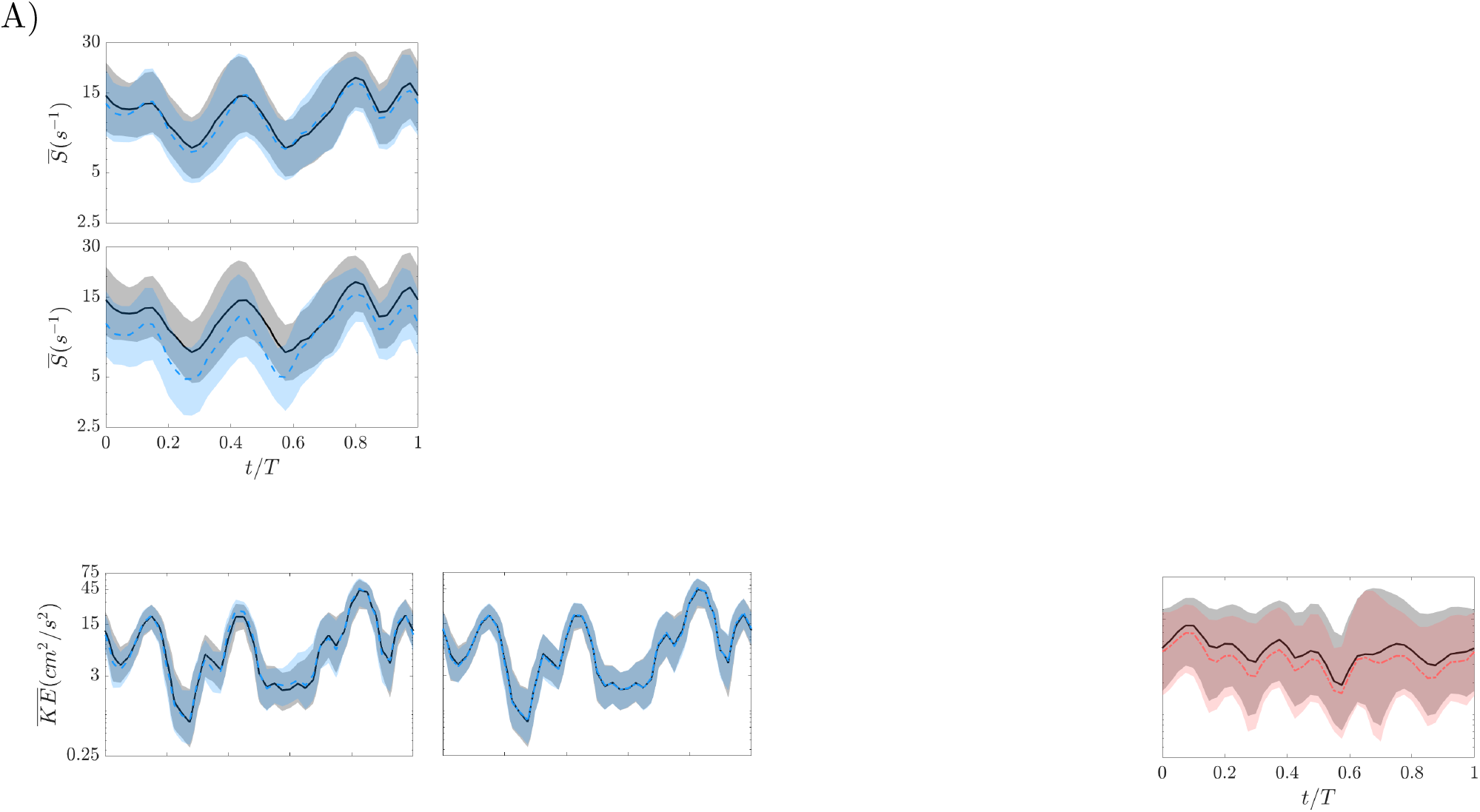
Shear rate and Kinetic energy in the distal left atrial appendage (LAA) for Newtonian and non-Newtonian different constitutive laws. **A)** Time histories during one cardiac cycle of shear rate median and 25^th^-75^th^ percentiles (denoted by colored shaded regions). The subject with normal atrial function (case 2) is represented with blue lines and patches, and the subject with an LAA thrombus and abnormal atrial function (case 5) is depicted in red. The four non-Newtonian constitutive laws are included for each case: hematocrit changes between rows, and the constitutive models between columns. Median and *25^th^-75^th^* percentiles of Newtonian case for the same patient are also included in black for comparison. **B)** Time histories during one cardiac cycle of kinetic energy median and 25^th^-75^th^ percentiles. Panels are presented in the same format than in **A)**.

## 4 Discussion

The left atrial appendage (LAA) is regarded as the most likely thrombosis site in patients with AF who suffer embolic strokes [11]. We hypothesized that sustained low shear leading to increased blood viscosity could cause hemodynamics alterations and magnify LAA blood stasis [27, 28], a critical factor in thrombogenesis [48]. This study aimed to evaluate how non-Newtonian rheology could affect left atrial (LA) hemodynamics and CFD-derived metrics of LAA blood stasis, particularly residence time. Since Shettigar *et al*. [49] reported blood residence time to examine the washout of artificial ventricles, this quantity has been used to study the risk of intracardiac thrombosis [12, 28, 50]. Patient-specific computational fluid dynamics (CFD) analyses are emerging as a tool to help assess LAA thrombosis risk in AF patients [23] and after LAA occluder device implantation [21]. However, previous investigations have considered constant fluid viscosity, neglecting the non-Newtonian rheology of blood at low shear rates.

### 4.1 Red blood cell aggregation in the LA and LAA

Red blood cell (RBC) aggregation into rouleaux is a significant cause of non-Newtonian blood rheology at low shear rates and long residence times. There is ample clinical evidence of RBC aggregation in the left atrium because RBC rouleaux increase blood’s echogenicity creating spontaneous contrast (a.k.a. “smoke”), which are readily detected in transesophageal ultrasound images [29]. Multiple studies have established a relationship between LAA blood stasis, spontaneous echocardiographic contrast, and the risk of LAA thrombosis, especially in patients with AF [48, 30]. Furthermore, previous CFD analyses of LA hemodynamics consistently report shear rates below 70 s^−1^ and residence times above 5 s inside the LAA [27, 28, 51], which, according to data from *in vitro* experiments, should cause RBC aggregation. However, there is a lack of studies evaluating the importance of non-Newtonian blood rheology in the LA and LAA hemodynamics. Recently, Wang *et al*. [52] incorporated non-Newtonian effects in their CFD simulations of a patient-specific LA model to investigate how cardiac rhythm affects the risk of thrombosis. Since non-Newtonian effects were not Wang et al.’s [52] focus, they did not present Newtonian simulation results or explored different *Hct* values, making it difficult to compare their results with the present study.

### 4.2 The range of Non-Newtonian effects: impact of hematocrit and rouleaux formation kinetics

We considered constitutive relations based on the Carreau-Yasuda model (eq. 3), which has been extensively used in previous CFD analyses of cardiovascular flows outside of the LA [53, 54, 40]. The Carreau Yasuda model depends on several parameters that are sensitive to the hematocrit level. Hematocrit is a major determinant of blood viscosity together with flow shear rate [24]. Normal hematocrit levels can vary among individuals within the range *Hct* = [34, 54] depending on factors like age and sex [44, 45]. Thus, to evaluate the extent of potential non-Newtonian effects, we parameterized the Carreau-Yasuda model using values from the literature corresponding to *Hct* = 37 and *Hct* = 55 [41, 42]. These simulations, denoted CY-37 and CY-55, allowed us to probe non-Newtonian effects for the lower and upper limits of the normal *Hct* range. They yielded equal or higher blood viscosity inside the LA chamber as compared to the Newtonian simulations. The differences were especially relevant inside the LAA, where the normalized average viscosity (i.e., 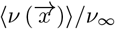) ranged between [1.18–1.38] for *Hct* = 37 and [1.98–2.56] for *Hct* = 55. These viscosity amplifications are consistent with those reported in CFD analyses of other vessels and cardiac chambers [40, 54]. Interestingly, despite the differences in 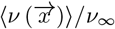 caused by changes in *Hct*, the non-Newtonian effects on the LAA flow patterns were appreciable for both the CY-37 and CY-55 constitutive relations (Figures 3–4, Figures SI 3-SI 6, Table SI 1). A possible explanation for this similarity could come by expanding the viscous stresses in the Navier-Stokes equation as 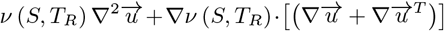, where the first term is proportional to the viscosity value and the second one is proportional to viscosity gradients. Thus, it is conceivable for non-Newtonian effects to be significant not only in regions with increased viscosity but also in regions with viscosity gradients, like the LAA of the CY-37 cases (see Figure 6A).

The Carreau-Yasuda model presumes that rouleaux formation exclusively depends on shear rate, which is a reasonable assumption for quasi-steady flows in small vessels with simple tubular geometries (*e.g*., the microcirculation). In these geometries, the flow timescales are longer than those of RBC aggregation kinetics, and low shear rate implies flow separation and long residence time. However, these assumptions break down for the complex, unsteady flows observed in the cardiac chambers [55]. *In vitro* investigations of rouleaux formation kinetics have reported RBC aggregation times between 1 and 6 s for normal blood and between 0.4 and 3 s for pathologically hypercoagulable blood [46, 47]. To consider the interplay between these timescales and the flow timescales induced by cardac pulsatility, we ran simulations with a thixotropic constitutive relation. Specifically, we implemented a modified constitutive relation that progressively activates the Carreau-Yasuda model as residence time increases beyond *T_μ_* = 3 s (eqs. 4–5) [25]. We considered hematocrit levels *Hct* = 37 and *Hct* = 55, and denoted the corresponding simulations CY-*T_R_*-37 and CY-*T_R_*-55. These simulations produced viscosities close to the Newtonian reference value inside the LA body, which is replenished every 1-2 heartbeats. However, the LAA remained a site of significant non-Newtonian rheology, showing normalized average viscosities of up to 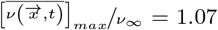 and 2.16 in the CY-*T_R_*-37 and CY-*T_R_*-55 cases considered, respectively. Our observations in the LAA contrast with the finding that considering the kinetics of rouleaux formation notably decreased shear-thinning effects in abdominal aortic and intracranial aneurysms [25]. We attribute this difference to the fact that blood residence time inside the LAA seems to be considerably longer than in aneurysms [25, 56].

### 4.3 Effects of non-Newtonian rheology on LA and LAA hemodynamics and blood residence time

There is scarce information about the influence of blood’s shear-thinning rheology on intracardiac hemodynamics. In the LV, clinical observation of spontaneous echocardiographic contrast suggest RBC aggregates can form in this chamber [29], and two modeling studies have evaluated non-Newtonian effects. Doost *et al*. [54] used CFD analysis to evaluate various constitutive relations on a patient-specific LV anatomical model, finding that blood viscosity increased enormously in the apical region. More recently, Riva *et al*. [57] analyzed patient-specific 4-D flow MRI and *Hct* measurements, reporting significant deviations from Newtonian behavior in areas of recirculating or low velocity. It should be noted that Doost et al.’s constitutive models correspond to high hematocrit (e.g., their Carreau model is equivalent to the present CY-55 model). Moreover, both Doost et al. and Riva et al. neglected the kinetics of rouleaux formation. Yet, blood flow tends to be slowest and residence time tends to be highest near the LV apex [58], and these features are exacerbated after acute myocardial infarction [12]. Overall, the existing data suggest that non-Newtonian effects in the LV might be appreciable in some pathological conditions.

The influence of blood’s shear-thinning on atrial hemodynamics is even more understudied than in the LV; we are unaware of previous flow imaging studies, and we are only aware of the beforementioned CFD study of Wang et al. [52]. Inside the atrial body, our Newtonian and non-Newtonian simulations produced similar flow patterns for all the constitutive models considered, even if viscosity significantly increased throughout the whole chamber for the CY-55 model. Likewise, TR inside the LA body was not appreciably affected by blood’s shear thinning. One can explain this behavior by considering that the mean flow pattern in the LA body consists of approximately straight motion from the PVs to the mitral annulus, without large-scale recirculation cells. The overall flow pattern is mainly dictated by global mass conservation, driving a fluid volume equal to the LV stroke volume (LVSV) through the LA body every heartbeat. Consequently, the mean *T_R_* inside the LA body could be roughly approximated as *T_R,LA-body_* ≈ *LAV*/ (*αLVSV*) s, where LAV is the mean LA volume and the factor *α* > 1 in the denominator accounts for most of the transport occurring during the E wave, which only spans a fraction of the cycle. While this estimation neglects relevant details such as the E/A ratio or the LA’s reservoir and booster functions, Figure SI 2 indicates that it is a good approximation for *α* ≈ 2, supporting that *T_R_* in the LA body is indeed dictated by global chamber mass conservation, at least in our simulations. Thus, since the LA volume, LVSV, and E/A ratio were the same in our Newtonian and non-Newtonian simulations, the overall distribution of TR in the atrial body was not significantly altered by implementing different constitutive laws.

In contrast to the LA body, the LAA is a closed chamber excluded from the atrium’s blood transit conduit that needs multiple heartbeats to be cleared and replenished. The filling and drainage jets driven by the expansion and contraction of the LAA walls play an important role in recycling the LAA’s blood pool. The net flow rate these jets create through the LAA’s orifice is dictated by the rate of change of LAA volume, which depends on the LAA contractility (and is hence reduced for AF subjects) but should be relatively insensitive to variations in blood viscosity. However, we and others have demonstrated the existence of significant secondary swirling flows inside the LAA [28, 59], which can also transport blood into and out of the LAA even if their associated net flux is zero. In fact, we previously showed that fixed-wall simulations and moving-wall simulations produce comparable values of blood residence time inside LAAs of the same geometry [28]. Furthermore, while previous CFD analyses indicate that LAA residence time correlates inversely with the LAA ejection fraction [28], this correlation is imperfect and additional factors such as LA volume, LAA morphology, PV flow profiles, and the MV have been associated to vortical patterns and residence time inside the LAA [60, 61, 21]. Of note, LAA morphology is a well-known stroke risk factor in patients with AF [11]. The LAA geometric factors that have been associated with stroke risk include left LAA trabeculations, orifice diameter [62], and lobe bend angle [63]. Our non-Newtonian simulations suggest that the blood shear thinning not only can affect the shape of the filling and draining jets but also dampen the secondary swirling motions inside the whole appendage. These differences translate into larger values of LAA residence time when non-Newtonian blood rheology is considered, especially in the distal region near the LAA apex.

Considering different constitutive laws and hematocrit values proved helpful to evaluate the influence of non-Newtonian effects in LAA residence time. In the cases studied, the time-averaged LAA TR increased by up to 38%, and and the maximum value of TR increased by up to 67%. Furthermore, significantly larger volumes within the LAA presented high values of *T_R_* when non-Newtonian effects were considered (*e.g*., the volume with *T_R_* > 6 s increased by up to 7-fold). Overall, our CFD analyses suggest that non-Newtonian blood rheology can significantly impacts the predicted values of LAA residence time, even for hematocrit within the normal range. Moreover, our data suggest that LAA blood stasis can be exacerbated by high hematocrit, consistent with clinically observed associations between elevated hematocrit and ischemic stroke risk, especially in males and the aged [64, 65]. Thus, while the current paradigm to explain this association focuses exclusively on alterations of cerebral blood flow [31], the present study implies that elevated hematocrit could also accentuate the risk of cardiac thromboembolism. Further studies are warranted to explore this potentially new mechanism.

### 4.4 Study Limitations

As mentioned above, the size of our subject cohort was too small (*N* = 6) to statistically infer or discard potential correlations between non-Newtonian effects and LA/LAA function or geometry.

The cohort was not designed to verify whether non-Newtonian effects exacerbate thrombus formation in AF patients because the patients with LAA thrombi or history of TIAs (cases 4-6) had persistent AF while the thrombus negative patients (cases 1-3) were in sinus rhythm. The primary objective of our study was to assess whether non-Newtonian effects can be significant inside the LAA in a wide range of conditions. With this objective in mind, we included subjects with normal LA function and a variety of LA dysfunctions (*e.g*., reservoir and/or booster pump) as well as subjects in sinus rhythm and with AF. Our results indicate that, in this diverse cohort, the inter-subject differences in blood stasis with Newtonian flow can be comparable to the intra-subject differences between Newtonian and non-Newtonian simulations. These analyses demonstrate the potential impact of non-Newtonian blood rheology in LA hemodynamics and LAA residence time.

We did not input patient-specific values of heart rate in our simulations. Instead, we ran all our cases at 60 beats per minute, similar to our previous constant viscosity simulations [28], which we used as the Newtonian baseline in the present study. Heart rate is known to influence both shear rate and residence time in the left ventricle [66, 67], and similar dependencies are anticipated in the LA. Future studies shall address whether these phenomena cause non-Newtonian effects in the LAA to be heart-rate dependent. We did not consider patient-specific hematocrit values. Instead, we ran simulations for two hematocrit values near the lower and upper limits of this parameter’s normal range to assess possible non-Newtonian effects. Likewise, the characteristic time (*T_μ_*) in our thixotropic constitutive laws was not patient-specific but equal to 3 seconds for all subjects based on human RBC aggregation times measured *in vitro* by different techniques [47, 46].

We implemented patient-specific inflow/outflow boundary conditions, with the caveat that the PV flow rates were evenly distributed to match the boundary conditions used in Garcia-Villalba et al.’s [28] Newtonian baseline simulations. Consequently, the resulting PV inflow profiles (blue curves in Figures 3–5) contained more peaks and valleys than the waveforms typically observed by, *e.g*., pulsed Doppler ultrasound [68]. While it is possible to derive a 0D model of the pulmonary circulation to determine each PV’s split of the total PV flow rate, fixing the value of the model impedances would be equivalent to directly specifying the flow split. The actual flow split is patient-specific and, given that the right lung has one more lobe than the left lung, the right PVs typically discharge more flow into the LA than the left PVs [22]. However, Lantz *et al*. [22] reported that flow velocities from simulations with evenly split PV flow rates agree reasonably well with patient-specific phase-contrast MRI data. Finally, we note that we did not model the geometry or motion of the mitral valve leaflets, based on previous evidence that they do not affect LA hemodynamics [15].

The calculation of the additional terms representing non-Newtonian effects in the Navier-Stokes equations had an insignificant impact on the overall computational cost per time step of the simulation. The most significant factor in comparing the cost of Newtonian and non-Newtonian simulations was varying the time step Δ*t* to keep the non-Newtonian *CFL* number below 0.2 across all timesteps of all simulations. As outlined in section 2.2, the non-Newtonian *CFL* has an additional term, i.e., *CFL_NN_* = *CFL_N_* +|∇ (Δ*ν_exp_*)|Δ*t*/Δ*x* that can grow in regions with strong viscosity gradients. In the CY simulations, *CFL_NN_* grew in regions with high velocity and low shear like the pulmonary veins, where both *CFL_N_* and |∇ (Δ*ν_exp_*)|Δ*t*/Δ*x* reached significant values. In these simulations, we found that decreasing Δ*t* by a factor of two with respect to the Newtonian simulations was sufficient to keep *CFL_NN_* < 0.2 in the whole domain. Thus, the CY simulations are approximately twice as expensive as the Newtonian ones. However, in the more realistic CY-TR simulations, non-Newtonian effects were not active in high-velocity regions regardless of the shear rate because these regions had residence time < *T_μ_*. Consequently, we were able to keep *CFL_NN_* < 0.2 in the CY-TR simulations using the same Δ*t* as the Newtonian ones. In short, the cost of adding realistic non-Newtonian effects on the simulations was negligible.

Our simulations are driven by mass flux boundary conditions so that a fluid volume equal to the patient-specific LVSV transits the atrium every heartbeat. This choice implies that the external work performed by the LA varies when the blood’s constitutive relation is varied for each subject. However, the pressure drop between the PVs and the MV changes less than 0. 3 mmHg between our Newtonian and non-Newtonian simulations, which is negligible compared to the total pressure in the LA (10 mmHg, see [69, 70, 71].

## 5 Conclusions

Patient-specific CFD analyses of left atrial hemodynamics suggest that the thixotropic, shear-thinning rheology of blood can significantly affect flow patterns inside the atrial appendage in a hematocrit-dependent manner. These non-Newtonian effects, which were neglected in previous CFD studies, aggravate blood stasis and increase the likelihood of left atrial appendage thrombosis, a recognized risk factor in ischemic stroke. Our simulations open a new possibility to untangle the contribution of hematocrit to stroke risk.

## Acknowledgements

This work was supported by grants from the American Heart Association (20POST35200401), the UCSD Galvanizing Engineering and Medicine Program, the National Institutes of Health (1R01HL160024), the Spanish Agency of Research (PID2019-107279RB-I00 and AEI/10.13039/501100011033), and the Regional Government of Madrid, Spain (Y2018/BIO-4858 PREFI-CM). A reciprocal visiting chair funded by the UC3M-Santander Foundation and computational time provided by XSEDE (Comet) and RES (Altamira and Caléndula) are gratefully acknowledged. Scanning electron microscopic images of RBC rouleaux (https://phil.cdc.gov/Details.aspx?pid=10902) used in the Graphical Table of Contents is also gratefully acknowledge.

