## Supporting information for "Non-Newtonian Blood Rheology Impacts Left Atrial Stasis in Patient-Specific Simulations"

### Supporting information (SI)

#### Non-Newtonian model validation

The benchmark configuration selected for validating the non-Newtonian model in TUCAN was extracted from Griffiths 2020 [1]. It consists of flow between two parallel walls driven by a constant pressure gradient in the streamwise direction, $-\rho^{-1}\partial p/\partial x = G$, where $p$ is the fluid pressure, $\rho$ is the constant density of the fluid and $x$ is the streamwise direction. We used the non-Newtonian constitutive law for the kinematic viscosity based on the Carreau-Yasuda eq. **??** in the main text of the manuscript. The parameters of the model were selected to compare with the analytical solutions of Griffiths, with $\nu_\infty \nu_0 = 1.25\times10^{-4}$ and $a = 2$. To explore the robustness of the code for different shear-thinning fluids $\lambda$ and $n$ were varied between 1/2 and 3/2 and between 1/3 to 2/3, respectively as in Griffiths 2020 **??**[1] (see FIG. 2 in ref. **??**). A simulation with the values of $n = 1/3$ and $\lambda = 3.313$ corresponding to our high-hematocrit $Hct = 55$ cases was also compared with the analytical solution. The simulations were 3D with periodic boundary condition in the directions parallel to the walls ($x$, and $z$), and a Dirichlet boundary conditions equal to 0 on the walls ($y = -h$, and $y = +h$).

For the selected values of $n$ and $\lambda$ the solution of the problem corresponds to a laminar steady streamwise velocity profile, $u\,(y)$, uniform in the wall-parallel directions. In SI 1 this streamwise velocity profile is shown for all cases. The profiles obtained with the non-Newtonian model implemented in TUCAN (solid lines) are compared with the analytical solutions from Griffiths (solid circles), showing agreement for all values of $\lambda$ and $n$ considered. The maximum differences of $u\,(y)$, are observed in the case with $n = 1/3$ and $\lambda = 3.313$ yielding a maximum relative error of 1.9%. The rest of the cases studied show relative errors in streamwise velocity lower than 1%.

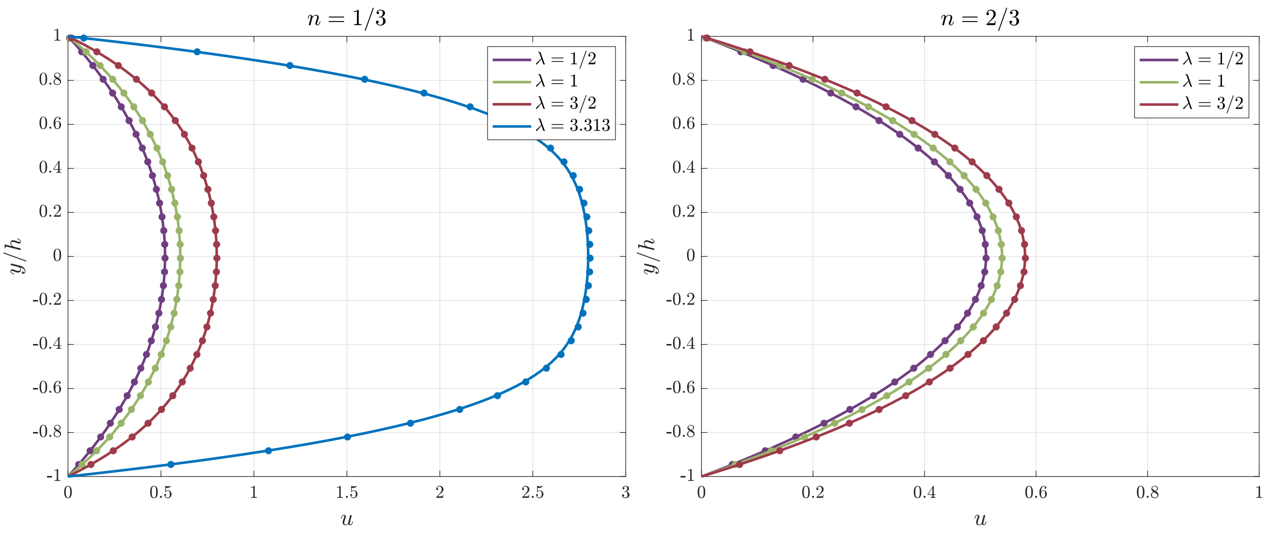


Figure SI 1: **Validation of modified version of TUCAN accounting for non-Newtonian effects** Streamwise velocity profile ($u$) along the vertical direction obtained with CFD simulations performed with the modified version of TUCAN accounting for non-Newtonian effect (solid lines), and analytical solutions from Griffiths 2020 [1] (solid circles).

#### Additional Figures and Tables

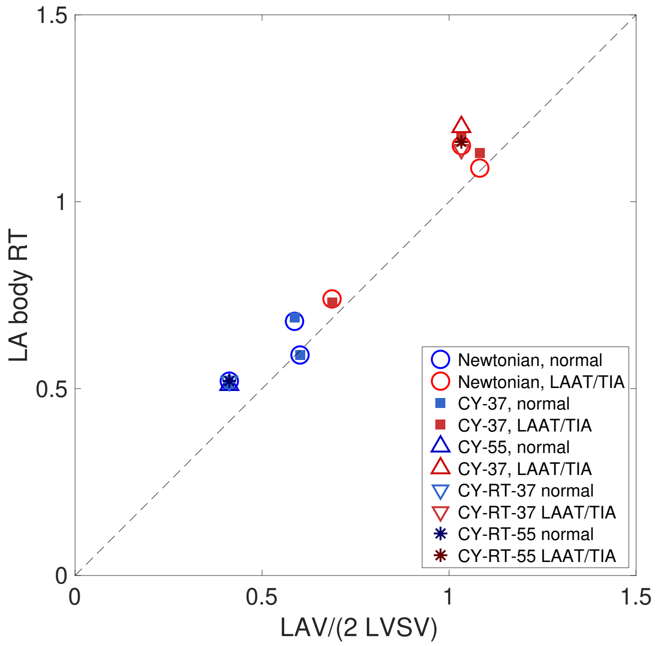


Figure SI 2: **Model to predict $T_R$ inside of the LA body.** Residence time ($T_R$) approximation based on the mean LA volume (LAV), the LV stroke volume (LVSV), and a factor $\alpha$ that accounts for the fraction of the cycle the E wave spans. Symbols identify the average value obtained for subjects with normal LA function (normal) cases and subjects with LAA thrombus or a history of TIAs (LAAT/TIA) with each viscosity model used (see Table **??**).

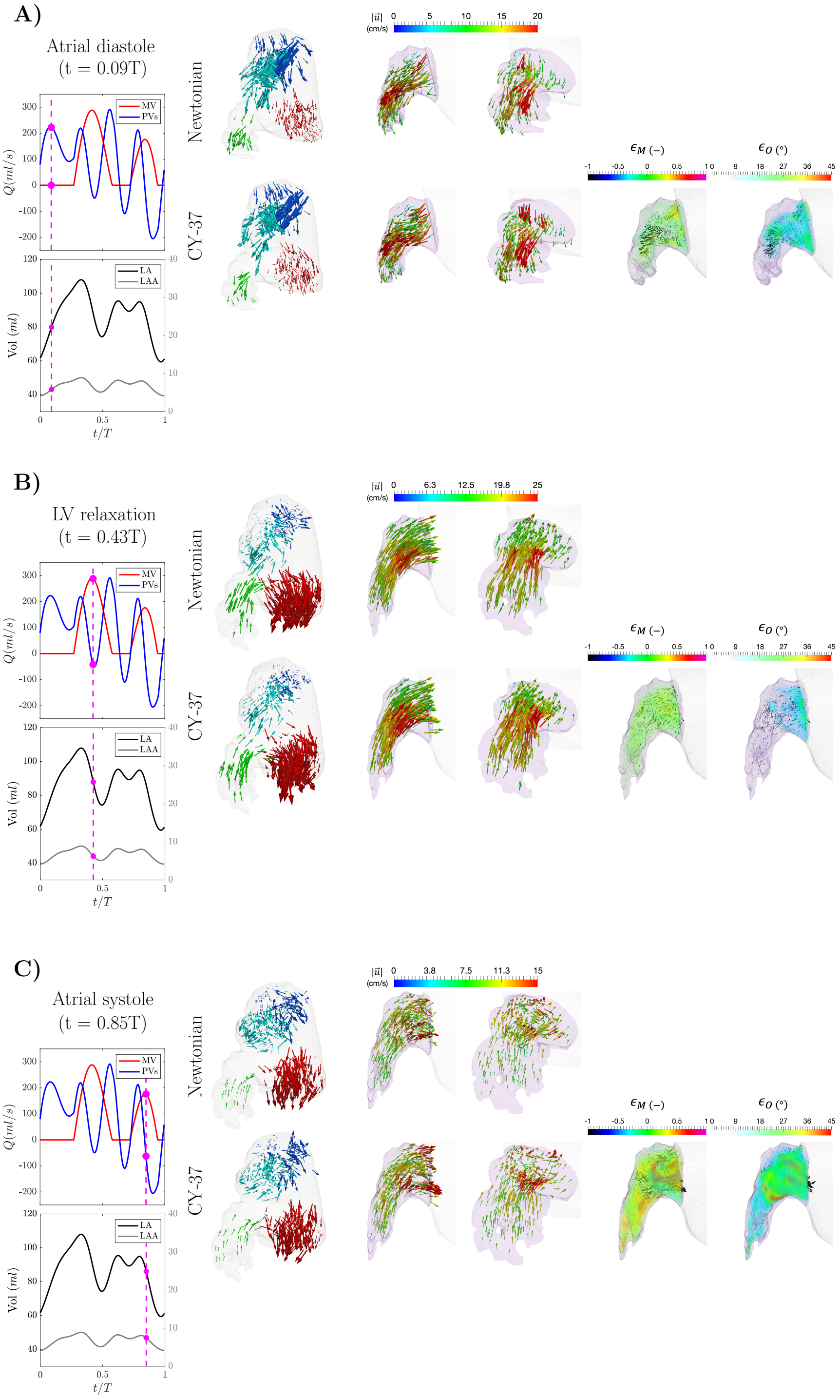


Figure SI 3: **Flow visualization of left atrial and left atrial appendage (LAA) hemodynamics from Newtonian and non-Newtonian simulations. Case1: Subject with normal atrial function and no LAA thrombus.** Vector maps of the 3-D blood flow velocity in the whole left atrium ($1^{st}$ column), two amplified views of the LAA in different orientations ($2^{nd}$ and $3^{rd}$ columns), and two LAA views showing the differences in velocity magnitude ($4^{th}$ column) and orientation ($5^{th}$ column) between Newtonian and non-Newtonian flow, using the same format as Figure **??**. **A)** Atrial diastole and peak flow rate through the pulmonary veins (t = 0.09 s). **B)** Left ventricular diastole and peak flow rate through the mitral valve (E-wave, t = 0.43 s). **C)** Atrial systole and peak backflow rate through the pulmonary veins (t = 0.85 s).

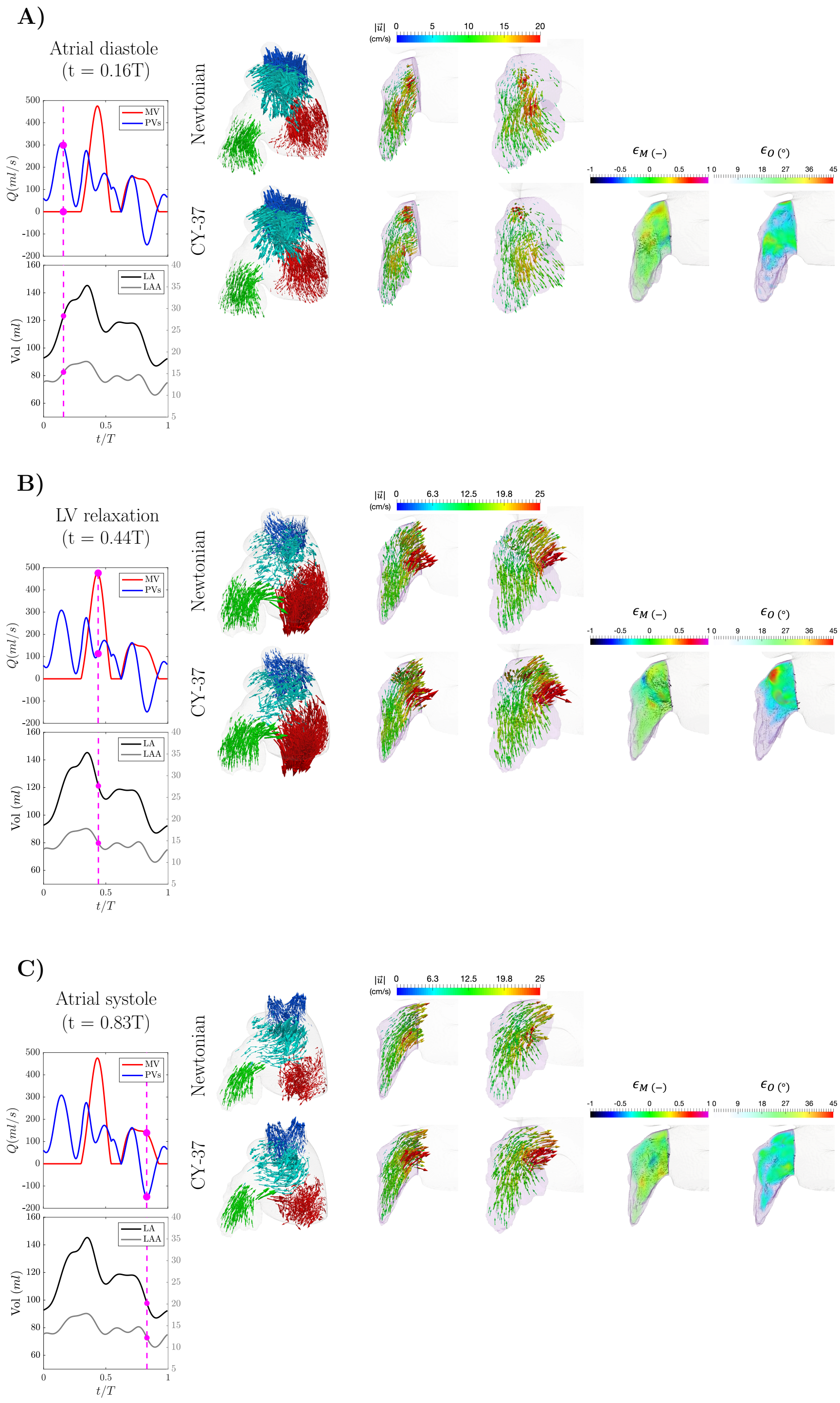


Figure SI 4: **Flow visualization of left atrial and left atrial appendage (LAA) hemodynamics from Newtonian and non-Newtonian simulations. Case3: Subject with normal atrial function and no LAA thrombus.** Vector maps of the 3-D blood flow velocity in the whole left atrium ($1^{st}$ column), two amplified views of the LAA in different orientations ($2^{nd}$ and $3^{rd}$ columns), and two LAA views showing the differences in velocity magnitude ($4^{th}$ column) and orientation ($5^{th}$ column) between Newtonian and non-Newtonian flow, using the same format as Figure **??**. **A)** Atrial diastole and peak flow rate through the pulmonary veins (t = 0.16 s). **B)** Left ventricular diastole and peak flow rate through the mitral valve (E-wave, t = 0.44 s). **C)** Atrial systole and peak backflow rate through the pulmonary veins (t = 0.83 s).

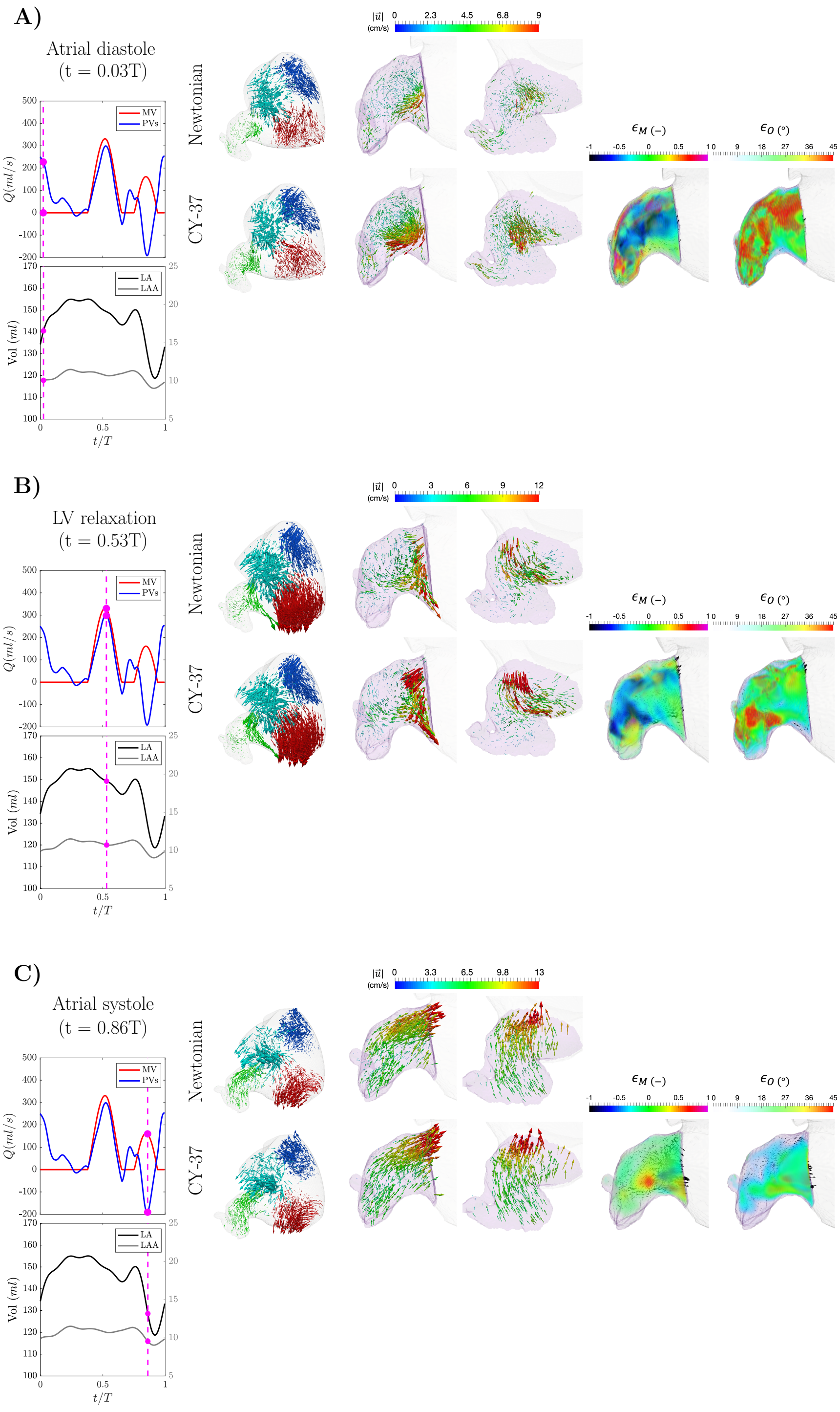


Figure SI 5: **Flow visualization of left atrial and left atrial appendage (LAA) hemodynamics from Newtonian and non-Newtonian simulations. Case 4: Atrial fibrillation patient with history of transient ischemic attacks (TIAs).** Vector maps of the 3-D blood flow velocity in the whole left atrium ($1^{st}$ column), two amplified views of the LAA in different orientations ($2^{nd}$ and $3^{rd}$ columns), and two LAA views showing the differences in velocity magnitude ($4^{th}$ column) and orientation ($5^{th}$ column) between Newtonian and non-Newtonian flow, using the same format as Figure **??**. **A)** Atrial diastole and peak flow rate through the pulmonary veins (t = 0.03 s). **B)** Left ventricular diastole and peak flow rate through the mitral valve (E-wave, t = 0.53 s). **C)** Atrial systole and peak backflow rate through the pulmonary veins (t = 0.86 s).

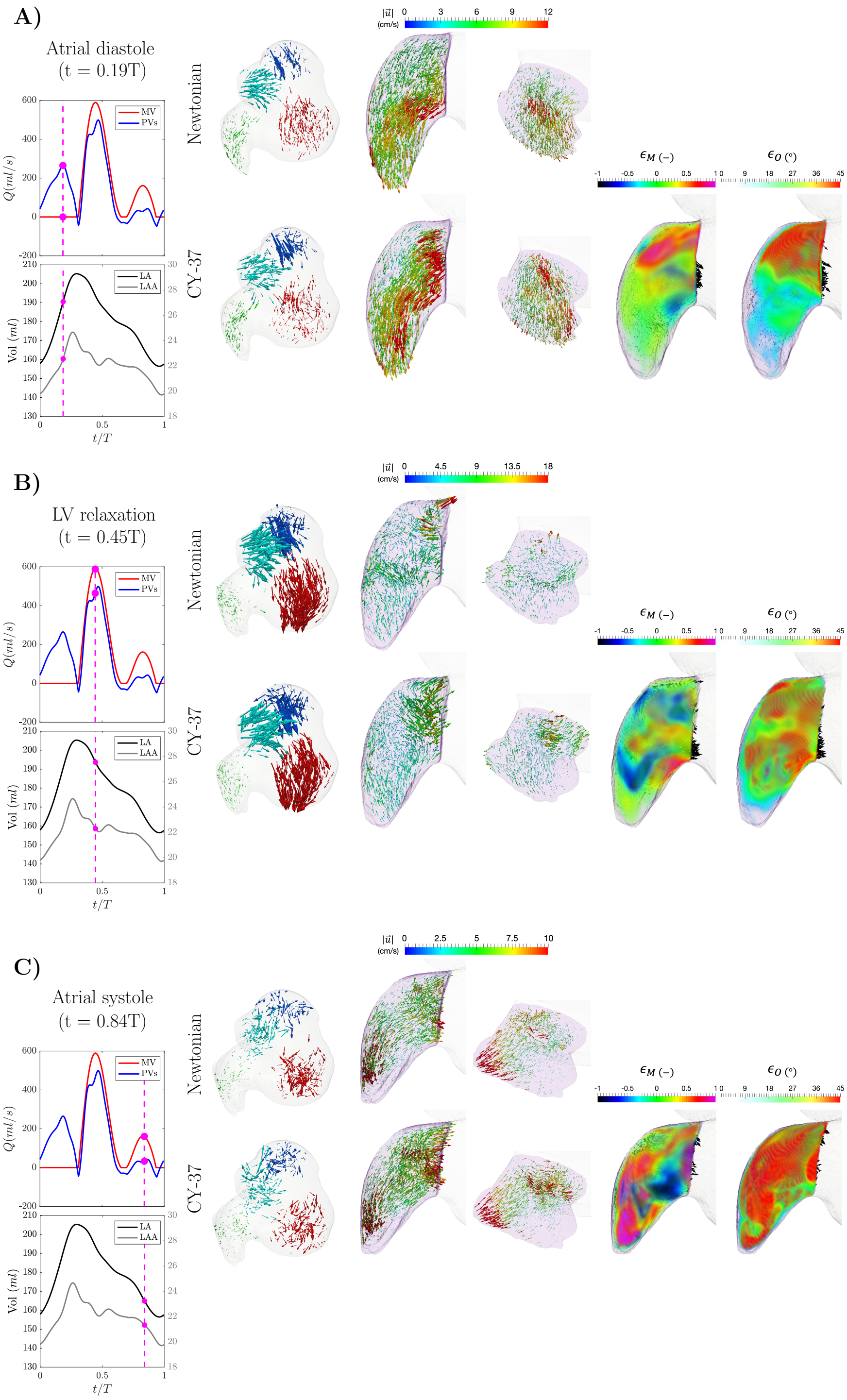


Figure SI 6: **Flow visualization of left atrial and left atrial appendage (LAA) hemodynamics from Newtonian and non-Newtonian simulations. Case 6: Atrial fibrillation patient with LAA thrombus (digitally removed before running the simulations).** Vector maps of the 3-D blood flow velocity in the whole left atrium ($1^{st}$ column), two amplified views of the LAA in different orientations ($2^{nd}$ and $3^{rd}$ columns), and two LAA views showing the differences in velocity magnitude ($4^{th}$ column) and orientation ($5^{th}$ column) between Newtonian and non-Newtonian flow, using the same format as Figure **??**. **A)** Atrial diastole and peak flow rate through the pulmonary veins (t = 0.19 s). **B)** Left ventricular diastole and peak flow rate through the mitral valve (E-wave, t = 0.45 s). **C)** Atrial systole and peak backflow rate through the pulmonary veins (t = 0.84 s).

| **Subject Number** | **Viscosity Model** | $\frac{1}{T}\int_0^T \left[\frac{1}{V_{LAA}}\int_V \epsilon_O dV\right] dt$ (°) | $\frac{1}{T}\int_0^T \left[\frac{1}{V_{LAA}}\int_V \lvert\epsilon_M\rvert dV\right] dt$ (−) |
|---|---|---|---|
| 1 | CY-37 | 16.3 | 0.126 |
| 2 | CY-37 | 24.8 | 0.184 |
| 2 | CY-$T_R$-37 | 26.9 | 0.198 |
| 2 | CY-55 | 24.4 | 0.179 |
| 2 | CY-$T_R$-55 | 22.2 | 0.162 |
| 3 | CY-37 | 23.7 | 0.202 |
| 4 | CY-37 | 23.2 | 0.221 |
| 5 | CY-37 | 21.3 | 0.152 |
| 5 | CY-$T_R$-37 | 21.1 | 0.172 |
| 5 | CY-55 | 19.9 | 0.164 |
| 5 | CY-$T_R$-55 | 26.9 | 0.229 |
| 6 | CY-37 | 41.8 | 0.296 |

Table SI 1: **Flow difference quantification inside the left atrial appendage (LAA) between non-Newtonian and Newtonian simulations.** The angular difference between velocity vectors, and the normalized difference of velocity magnitude of non-Newtonian ($\overrightarrow{u}^{NN}$) and Newtonian ($\overrightarrow{u}^{N}$) simulations are quantified using $\epsilon_O = \arccos\left(\frac{\overrightarrow{u}^{NN}\cdot\overrightarrow{u}^{N}}{||\overrightarrow{u}^{NN}||||\overrightarrow{u}^{N}||}\right)$ and $\epsilon_M = 2\frac{||\overrightarrow{u}^{NN}||-||\overrightarrow{u}^{N}||}{||\overrightarrow{u}^{NN}||+||\overrightarrow{u}^{N}||}$, respectively. The values of $\epsilon_O$ (in degrees) and $\epsilon_M$ presented in this table are integrated in the LAA during the whole cardiac cycle.
